# Diverse horizontally transferred cellulose biosynthesis gene clusters in *Escherichia coli* strains

**DOI:** 10.1101/2025.05.09.653004

**Authors:** Seyedmohammad Hosseinpourlamardi, Shaiqa Labiba, Li Li, Ali Dadvar, Ute Römling

## Abstract

The phosphoethanolamine modified exopolysaccharide cellulose is a major extracellular matrix component of *Escherichia coli* and *Salmonella typhimurium*. Upon enhanced acute virulence, however, cellulose production can be diminished or entirely abolished. Here, we report that homologs of the *bcsABC* core genes of the core genome *bcs* cellulose biosynthesis operon and even entire *bcs* operons can be mobilized on plasmids and are occasionally manifested on *E. coli* chromosomes at alternative locations. While BcsA2 and BcsA3 cellulose synthases are restricted to the genus *Escherichia*, the BcsA4 cellulose synthase and the entire associated *bcs* gene cluster are highly similar to one of the two chromosomal *bcs* gene clusters harboured by *Klebsiella pneumoniae*. Cyclic di-GMP turnover proteins known to post-translationally regulate cellulose biosynthesis are frequently localized on *bcs* operon bearing plasmids. In thermotolerant meat derived *E. coli* 740v1 harbouring a type 4 *bcs* operon on a plasmid, chemical and genetic evidence indicates that cellulose is produced in M9 minimal medium and upon activation by the second messenger cyclic di-GMP, an allosteric activator of the cellulose synthase, respectively. With gene duplication and horizontal gene transfer to contribute to the multiplication of *bcs* operons in a number of different bacterial species, the ecological role(s) of multiplication, transfer and replacement of cellulose biosynthesis operons also in species other than *E. coli* still need to be unraveled.

## Introduction

Poly 1,4-beta-D-glucan macrofibrills is an exopolysaccharide synthesized by bacteria, oocytes, animals and plants (Klemm et al. 2005, Fugelstad et al. 2009, Nakashima et al. 2004, Pedersen et al. 2023). With the linear glucan chains tightly assembled in parallel into micro- and macrofibrills by weak, but numerous hydrogen bonds, the chemically inert cellulose I macromolecule is formed in bacteria as investigated with *Komagataeibacter* spp. as major model organisms. Widespread in diverse branches of the bacterial domain of life, cellulose is a component of the extracellular matrix of biofilms, multicellular, predominantly sessile, assemblies of autonomous microbial cells displayed as cell aggregates, microbial communities on abiotic or biotic surfaces or pellicle formation at the air-liquid interface among other biofilm modes of growth (Romling and Galperin 2015). Thereby, the catalytic core unit of cellulose biosynthesis, the cellulose synthase BcsA (also known as CelA) is embedded in different genomic contexts with different accessory elements (Romling and Galperin 2015). For example, in Gram-negative bacteria, the *bcsA* cellulose synthase gene is colocates with *bcsB* encoding a periplasmic protein that interacts with a C-terminal transmembrane helix with BcsA and is essential for the catalytically activity of the cellulose synthase complex. Further *bcsC* codes for the outer membrane pore to release the glucan chain from the periplasm into the extracellular space. BcsZ coding for a type 8 glucanase is the forth gene of the core unit required for cellulose biosynthesis with a still not fully characterized role in biosynthesis and ecology (Ahmad et al. 2016, Verma et al. 2024, Robledo et al. 2012, Menendez et al. 2019) The currently experimentally verified cellulose synthase operons can be categorized into at least four different groups associated with different accessory genes. Group I comprises the cellulose biosynthesis gene cluster present in the fruit-rotting alpha-proteobacterium *Komagataeibacter xylinus* a model organism which has been shown to produce high amounts of unmodified poly-1,4 beta-glucan chains; the beta-proteobacterium *Burkholderia phymatum* and the gamma-proteobacterium *Dickeya dadantii*. Group I, II and III cellulose biosynthesis operons are present in Gram-negative bacteria and are characterized by the presence of BcsB. The tight functional coupling is indicated by the occasional observation of BcsAB fusion proteins. The bacterial cellulose macromolecule can be covalently modified by accessory gene products (Thongsomboon et al. 2018, Spiers et al. 2003). Thereby, group II cellulose biosynthesis operons are characterized by the *bcsEFG* accessory genes. Their gene products are required for optimal expression/integration of the cellulose synthase into the cytoplasmic membrane, synthesis of the beta-1,4-glucan chain and the covalent modification of the nascent glucan chain with phosphoethanolamine at the C6 of the glucose (Sun et al. 2018, Thongsomboon et al. 2018). Acetylation is another covalent modification of the 1,4-beta-glucan chain performed by gene products of the *wssFGHIJ* operon (Spiers et al. 2003). Despite possessing the same substrate specificity leading to the same product output, cellulose synthases and other gene products of the cellulose biosynthesis operon can display a high sequence diversity (Liu et al. 2020). On the other hand, recently, the first enzymes that synthesize mixed 1-3, 1-4 β-glucans have been identified in bacteria (Perez-Mendoza et al. 2015, Chang et al. 2023). Those enzymes are highly homologous to cellulose synthases.

The type IIa cellulose biosynthesis operon is highly conserved between *Escherichia coli* and *Salmonella typhimurium* with respect to gene conservation and chromosomal location (Zogaj et al. 2001). In these two species, cellulose, co-synthesized with equally conserved amyloid curli fimbriae, constitute the major components of the extracellular matrix of the rdar biofilm. Although rdar biofilm can be observed readily as a colony morphotype on agar plates under laboratory conditions, rdar biofilm permits environmental persistence and tolerance against stress factors. Thereby trades rdar biofilm formation, and in particular the biosynthesis of cellulose’, environmental survival against acute virulence including growth within macrophages (Pontes et al. 2015, Ahmad et al. 2016, Romling et al. 2000). Indeed, invasive pathovars of *E. coli* and *S. typhimurium*, such as *Shigella* spp., Enteroinvasive *E. coli*, other *E. coli* sequence types and invasive *S. typhimurium* ST313 have highly reduced or entirely lost the ability to synthesize cellulose (Nhu et al. 2024, Singletary et al. 2016).

Surprisingly though, in the case of *Shigella*, some of these pseudogenes have regained functionality under in vivo like conditions (Chanin et al. 2019). On the other hand, certain bacterial species such as *Komagataeinbacter* spp. which produces high amounts of cellulose possess more than one cellulose biosynthesis operon (Saxena and Brown 1995). This is especially surprising in the case of *Proteus sp*. which have not been reported to produce cellulose (Liu et al. 2020). Of note, those species possess only one copy of a GGDEF diguanylate cyclase producing the ubiquitous second messenger cyclic di-GMP which is hypothesized to post-translationally stimulate cellulose biosynthesis via the PilZ domain located at the C-terminal end of most bacterial cellulose synthases (Liu et al. 2020).

However, the role of cellulose biosynthesis in *E. coli* is more complex than previously anticipated. In this work, we report the introduction of alternative horizontally transferred cellulose biosynthesis operons coding for distinct cellulose synthases which contain a variable number of essential and accessory gene products, into *E. coli* strains. Thereby, a second cellulose biosynthesis gene cluster, which conventionally consists of the core components *bcsABC* was found to be plasmid or chromosomally encoded, while a type 4 *bcs* gene cluster introduced novel *bcs* cluster associated genes into *E. coli*. One of these plasmid encoded *bcs* gene clusters was activated by the second messenger cyclic di-GMP. While these observations indicate that genes that code for exopolysaccharide extracellular matrix components can be highly mobile even in human associated bacteria, the evolutionary forces that drive the dissemination and diversification of cellulose biosynthesis genes in *E. coli* including the thermotolerant meat-derived strain *E. coli* 730V1, and other bacterial species including plant symbionts still need to be unraveled.

## Results

### *Escherichia coli* strains can possess two BcsA cellulose synthases

Not only highly cellulose producing bacterial species like *Komagataeibacter* spp. possess more than one cellulose biosynthesis operon, even bacterial species/strains belonging to the families Morganellaceae and Enterobacteriaceae such as *Proteus* spp. and *Klebsiella* spp., respectively, possess more than one cellulose biosynthesis operon (Liu et al. 2020, Romling and Galperin 2015). Commonly, isolates of the species *E. coli* are considered to encode one conserved class IIa cellulose biosynthesis gene cluster consisting of the two divergently transcribed operons *bcsQRABZC* and *bcsEFG* on the core genome (Zogaj et al. 2001, Solano et al. 2002). *E. coli* strains, equally as the closely related *Salmonella* spp., have been observed, upon evolution, rather to loose the ability to synthesize cellulose (Nhu et al. 2024, Singletary et al. 2016). However, upon investigating the conservation of cellulose synthases in *E. coli* by BLAST screening for homologous proteins, we noted the occurrence of cellulose synthases only distantly related to the core genome BcsA1 cellulose synthase to be present in certain *E. coli* strains. We therefore set up to investigate the diversification and the prevalence of alternative cellulose biosynthesis gene clusters in *E. coli* in more detail.

A systematic analysis of the presence of distantly related BcsA cellulose synthases outside of the core genome class IIa cellulose biosynthesis operon in *E. coli* on the core genome and on plasmids uncovered three additional groups of cellulose synthases termed BcsA2-BcsA4 with distinct similarity to the core genome cellulose synthase BcsA1 (Figure 1A, S1A and B; Table S1). BcsA2 members show approx.. 43% identity and 57% similarity, BcsA3 members 41% identity and 55% similarity and BcsA4 members 35% identity and 50% similarity as compared to the core genome cellulose synthase BcsA1 (Figure 1A, S1A and B). We noted that the BcsA2 and BcsA3 cellulose synthase protein classes are not found outside of the *Escherichia* genus but occasionally observed in other *Escherichia* species (Blast analysis, Figure 1A and S1A). However, the BcsA4 type cellulose synthase, most distantly related to BcsA1, is most closely related to one of two cellulose synthases of *Klebsiella* pneumoniae MGH 7843 (Figure S1). BcsA2 to BcsA4 cellulose synthases, as the vastly investigated cellulose synthase of *Cereibacter sphaeroides* (Morgan, McNamara and Zimmer 2014), lack the extended approx. 150 amino acid long N-terminal end of *E. coli* BcsA1, but maintain the nine core transmembrane helices flanking the catalytic cytoplasmic domain, the signature amino acids indicative for processive type 2 glycosyltransferases, catalytic activity and cellulose synthase activity equally as the C-terminal PilZ domain required for cyclic di-GMP binding with the RxxxR and DxGxS binding motifs (Figure 1A, S1A and B). BcsA2-BcsA4, however, lack the extended C-terminus of approx. 30 amino acids. As concluded from these analyses, BcsA2 to BcsA4 cellulose synthases are predicted to be functional.

**Figure 1.**
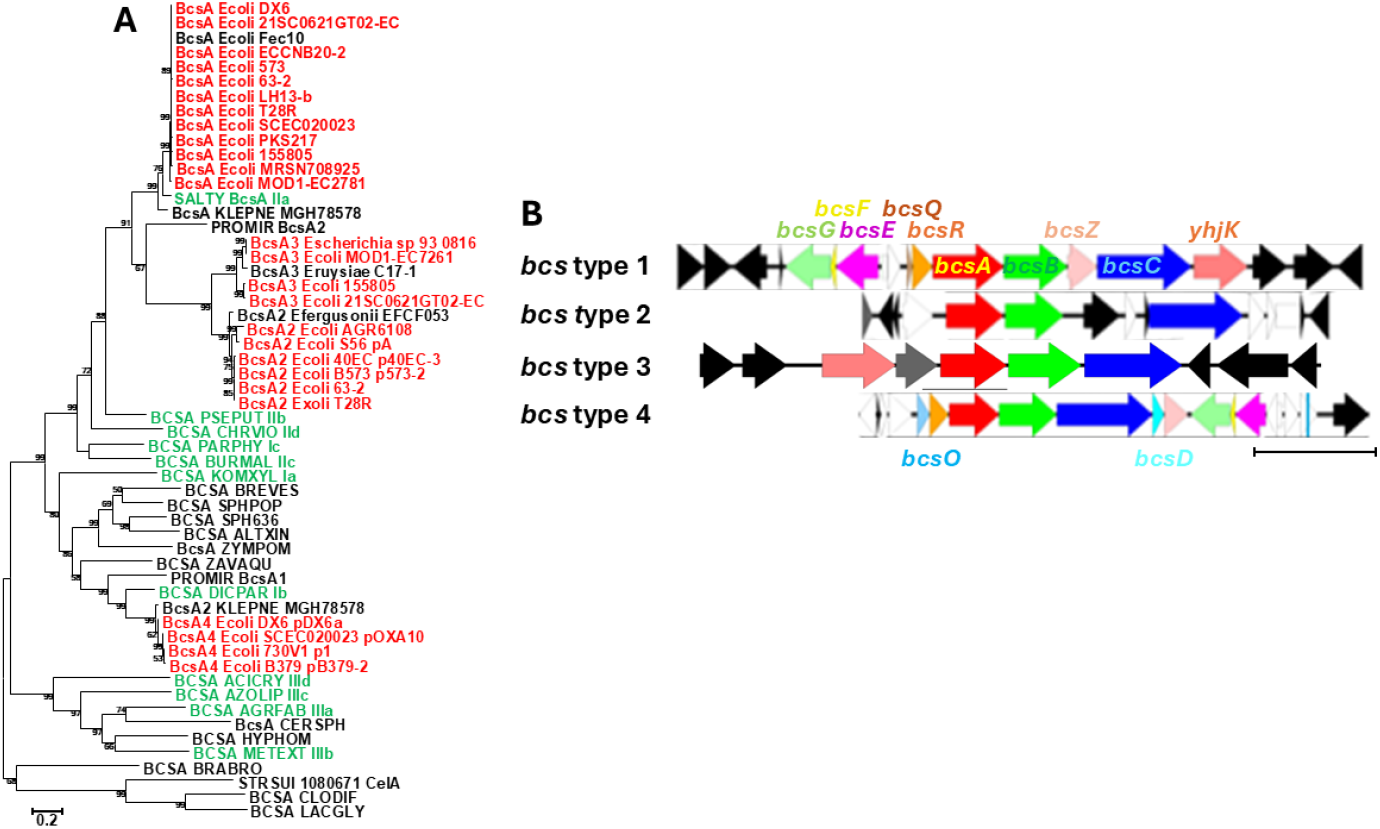
**1A**. Phylogenetic tree of representative *E. coli* cellulose synthases BcsA1-4 with reference BcsA proteins and genomic context of *bcsA1-4* genes. Selected BcsA cellulose synthases (see appendix and Figure S1A) were aligned with ClustalX and subsequently manually curated. The phylogenetic relationship has been assessed using a maximum likelihood (ML) based phylogenetic tree with 1000 bootstraps in MEGA 11.0. The nucleotide sequences and annotations of the *bcs* gene clusters and their gene neighborhoods were extracted from the genomic sequences stored at NCBI and aligned and visualized with Easyfig (Sullivan et al. 2011). *E. coli* and *Escherichia* BcsA sequences are shown in red and reference BcsA sequences (Romling and Galperin 2015) are shown in green. Size bar, substitutions per site. **1B**. Representative bacterial cellulose biosynthesis (*bcs*) gene clustersm of the four types; type 1, core genome *bcs* gene cluster encoding BcsA1 cellulose synthase and type 2-4, accessory *bcs* gene clusters encoding BcsA2-4 cellulose synthases. Designation of gene identity see Figure. White arrow, mobile element gene. Scale bar, 5 kbp.

### BcsA2, BcsA3 and BcsA4 are found in different genetic context

The three cellulose synthase classes BcsA2, BcsA3 and BcsA4 are integrated in different genetic contexts (Figure 1A and S2, Table S1). We observed that the BcsA2 cellulose synthase is in the majority of cases found embedded in a *bcsABC* gene cluster often located on plasmids >90 kbp in size with a p0111 origin of replication indicating phage origin of the plasmid (Pfeifer et al. 2021) and are on average larger than

*E. coli* plasmids without *bcsA* (Figure 1A; Figure S2 and S3), but can also be found exceptionally on the chromosome (for example *E. coli* strain T28R (Figure 2)). As far as it can be concluded from the available sequenced genomes and the genetic context on contigs *bcsA3* genes are embedded in a *bcsABC* gene cluster located on the chromosome. In the context of an extended *bcs* gene cluster, BcsA4 is encoded in the context of the *bcsOQABCDZ bcsEFG* gene cluster on large plasmids. The BcsA4 gene cluster is highly similar to a chromosomally encoded gene cluster from *Klebsiella* spp. (Figure S1).

**Figure 2.**
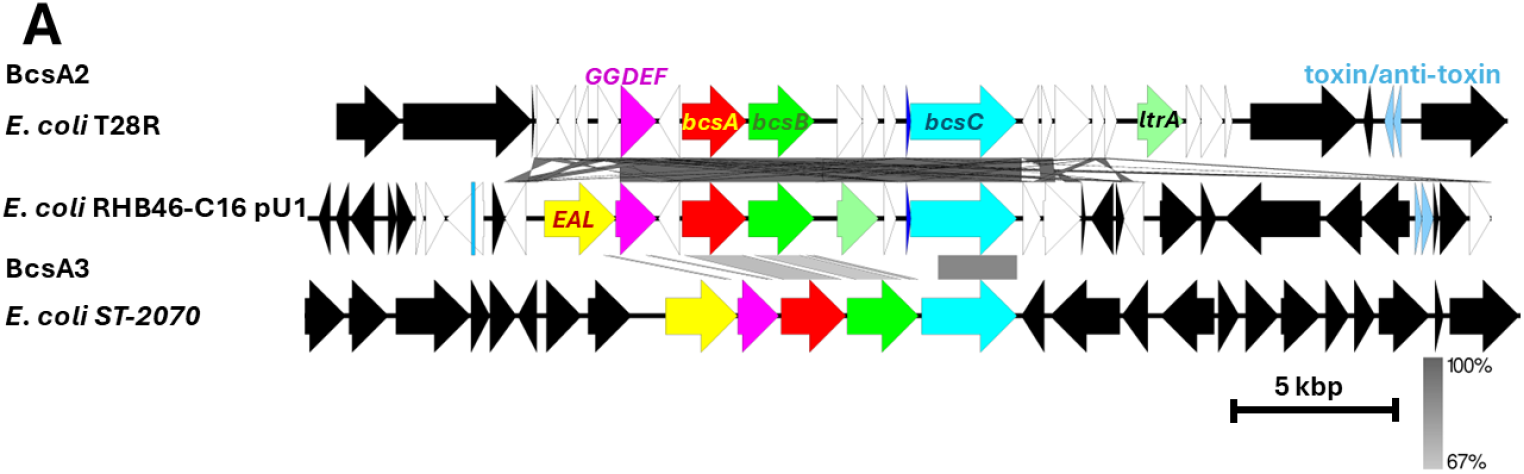
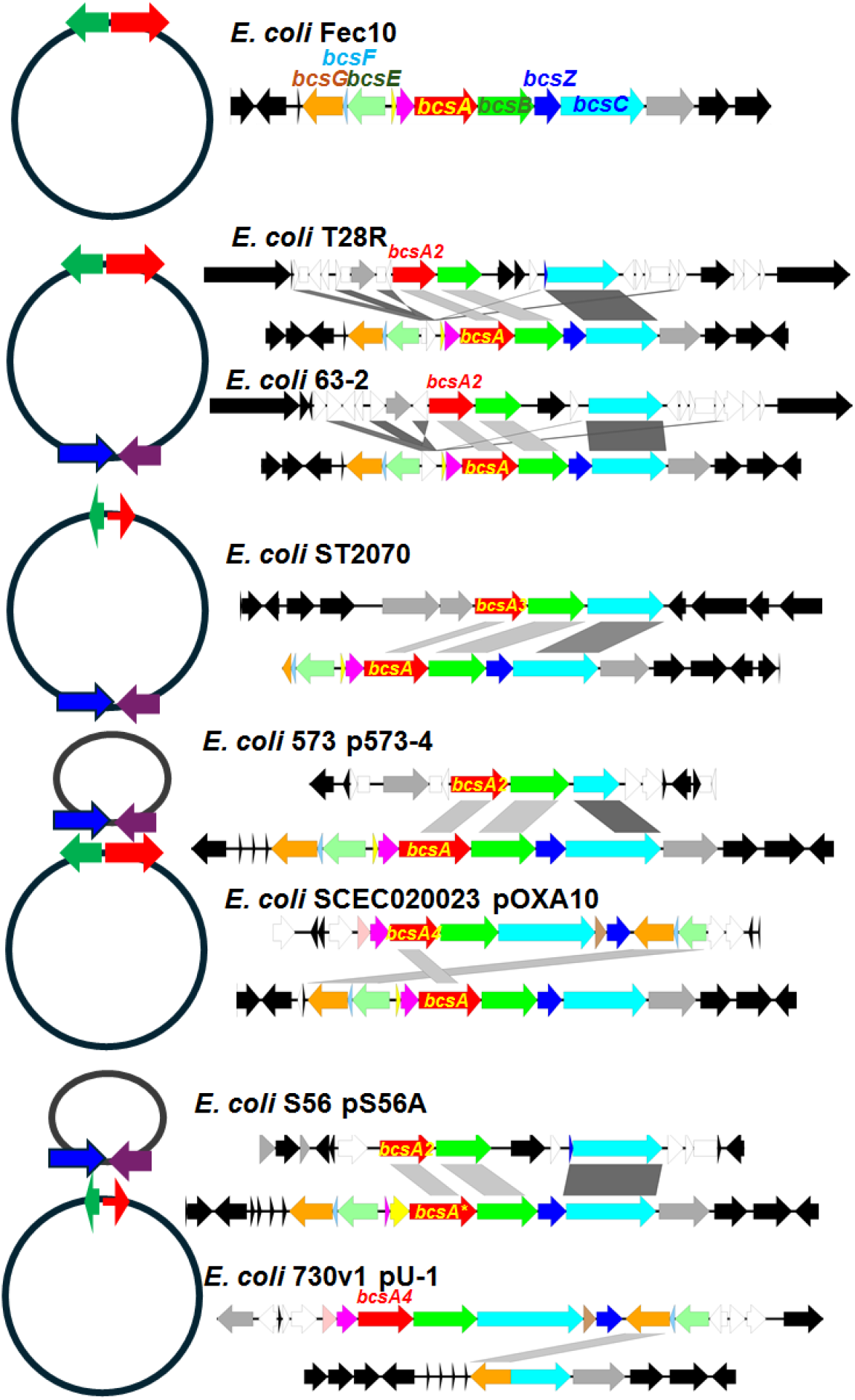
**2A**. Chromosomal BcsA2 and BcsA3 cellulose biosynthesis gene clusters. White arrow, mobile element gene. **2B**. Occurrence of *bcsA2-4* cellulose synthase operons in different contexts in *E. coli:* conventional; conventional + unconventional on chromosome; conventional and unconventional on plasmid; conventional deleted. Reference, *E. coli* Fec10 a modern commensal human ST10 isolate (Kamal et al. 2021) of *E. coli* K-12 MG1655. The nucleotide sequences and annotations of the *bcs* gene clusters and their gene neighborhoods were extracted from the genomic sequences stored at NCBI and aligned and visualized with Easyfig (Sullivan et al. 2011). Gray arrow, cyclic di-GMP turnover protein gene; white arrow, mobile element gene.

We subsequently investigated whether gene products associated with the BcsA cellulose synthases have a similar phylogenetic clustering which would indicate a common horizontal gene transfer or whether a mosaic structure of gene acquisition needs to be considered. To this end, we aligned and subsequently constructed phylogenetic trees of all the cellulose biosynthesis proteins that had been identified in the BcsA2-BcsA4 cellulose synthase gene clusters in the context of reference proteins (Figure S4). We found that BcsB required for catalytic activity of the cellulose synthase (and even in *E. coli* occasionally found as a fusion protein with BcsA (Figure S1A) and the outer membrane pore BcsC cluster concordantly as BcsA (Supplementary Figure S4A and B). Also the phylogenetic context of the cellulase BcsZ is congruentwith the phylogeny of BcsA (Figure S4C). BcsE and BcsG members of the core genome cellulose biosynthesis gene cluster are present in the context of the BcsA4 cellulose biosynthesis operon (Figure S4D and E). *BcsE4* encoding the GIL-type cytoplasmic c-di-GMP receptor required for optimal cellulose biosynthesis (Fang et al. 2014) has, however, consistently evolved as a pseudogene. BcsG4 members, required for stabilization of the cellulose synthase and covalent modification of the glucan chain (Sun et al. 2018, Thongsomboon et al. 2018) on the other hand built up their own cluster.

The accessory protein BcsD is located in the BcsA4-like gene cluster (Romling and Galperin 2015, Wong et al. 1990). BcsD a component of type I cellulose biosynthesis operons enhances the crystallinity of the cellulose macromolecule by interaction with the cellulose synthase complex and the nascent glucan chain (Figure S4F; (Sana et al. 2024)). The accessory protein BcsO is conventionally found in class Ib *bcs* gene clusters (Romling and Galperin 2015). Indeed, BcsO is infrequently found in cellulose biosynthesis operons. Proline-rich BcsO interacts in the cytoplasm with BcsD in *Dickeya dadantii* Ech703 and other plant-associated or environmental gamma-proteobacterial species (Figure S4G, (Sana et al. 2024)). We identified the closest homologue of BcsO by BLAST search. The phylogenetic tree of the evolutionary relationship showed that BcsO is highly similar to *K. pneumoniae* BcsO present in the second cellulose biosynthesis operon (Figure S4H).

### The Wss genes form a novel class of cellulose biosynthesis operons producing acetylated cellulose

Upon construction of the phylogenetic trees for the BcsA cellulose synthase and the periplasmic BcsB protein, we have included the representative cellulose synthases of class I to IV cellulose biosynthesis operons (Figure 1A, S1A, S1C). Assessment of the phylogenetic relationship between the cellulose synthases of the different classes and additional known and to be confirmed BcsA proteins indicates that, the enzymes that synthesize the macromolecule belong to a group of proteins with distinct similarity, while the glucan chains can be covalently modified by accessory proteins. Considering the Wss gene cluster which is characterized by accessory genes involved in the partial acetylation of various carbon positions at the glucose moieties (Spiers et al. 2003), we propose constitutes a new class Id cellulose biosynthesis gene cluster (Figure S1A). However, more distantly related potential cellulose synthases such as present in the alpha-proteobacterium *Zymomonas mobilis* (Cao et al. 2022, Jeon et al. 2012) and not categorized in any class exist. We also cannot exclude at this point that some of the proposed cellulose synthases such as CelA of *Streptococcus suis* are in fact mixed 1-3, 1-4 beta-glucan synthases (Figure S1C).

### Horizontal acquisition of a second cellulose biosynthesis operon results in different outcomes

In contrast to the gene cluster for poly-N-acetyl-glucoseamine biosynthesis, the chromosomal location of the conventional *bcsA1* cellulose biosynthesis operon is conserved (Cimdins et al. 2017). We were therefore wondering about the fait of the core genome *bcsA1* gene cluster upon acquisition of a second cellulose biosynthesis operon on the chromosome. Investigation of the reference strains harbouring an alternative cellulose biosynthesis operon showed that various constellations can occur (Figure 3). The newly introduced unconventional Bcs gene cluster can be either found on a large plasmid or on the chromosome. Thereby the location of the second *bcs* gene cluster on the chromosome is distinct from the *bcs1* gene cluster. Independently of the chromosomal or plasmid location of the horizontally transferred gene cluster, the conventional *bcs1* gene cluster is either present or has been deleted.

**Figure 3.**
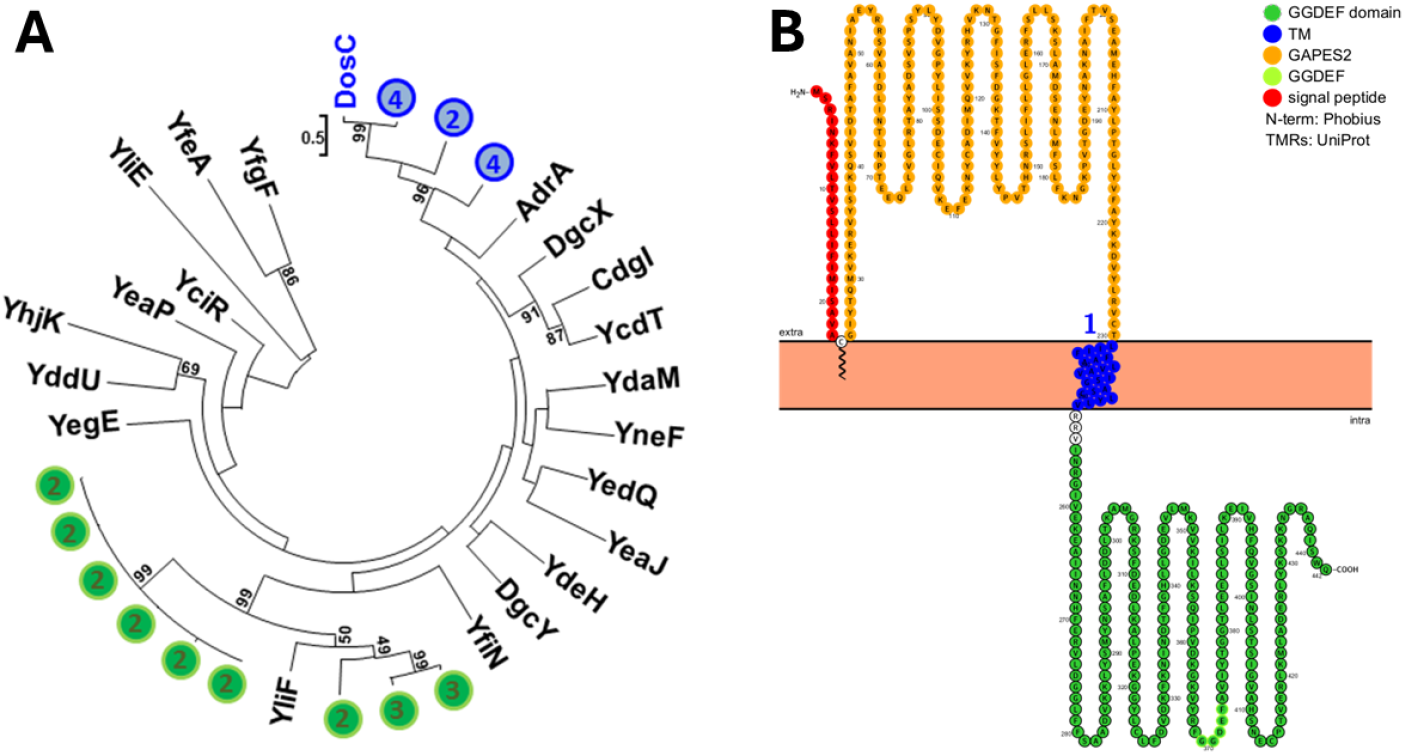
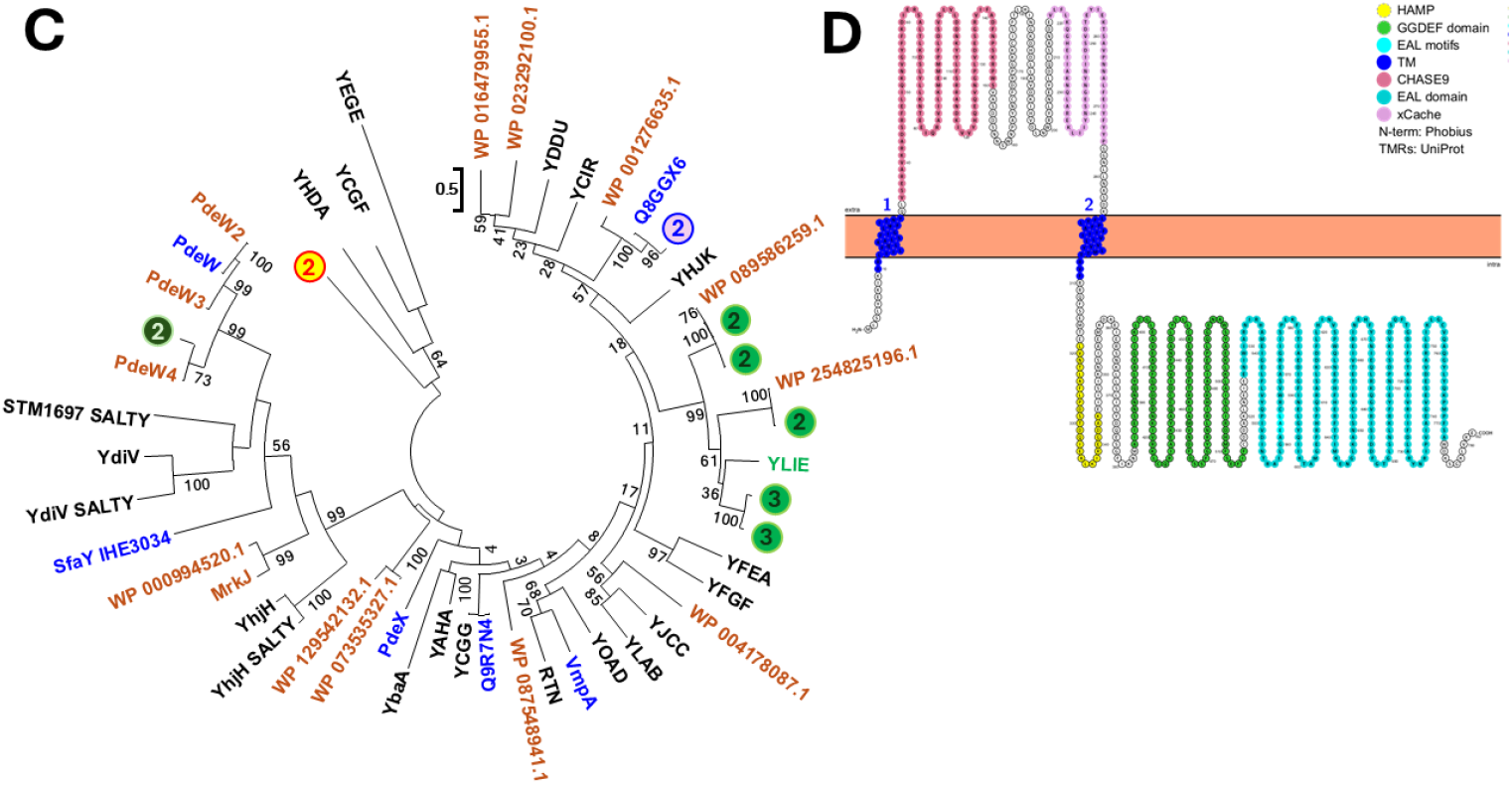
Phylogenetic tree to place GGDEF diguanylate cyclase and EAL phosphodiesterase domain proteins adjacent to *bcsA* type 2-4 cellulose biosynthesis gene clusters in the context of all cyclic di-GMP turnover proteins of *E. coli* MG1655 (identical with the modern ST10 strain *E. coli* Fec10 (Kamal et al. 2021)). **3A**. Phylogenetic tree of *E. coli* GGDEF domains of GGDEF domain proteins found adjacent to BcsA2-4 cellulose biosynthesis gene cluster(s) in the context of the GGDEF domains from chromosomally encoded proteins. GGDEF domain proteins adjacent to BcsA2-4 cellulose biosynthesis operons are homologous to YliF and DosC over the entire length of the amino acid sequence (Figure S5). Numbers indicate the BcsA cellulose synthase class adjacent to the GGDEF domain protein. **3B**. Protter image of the chromosomally encoded GGDEF domain protein YilF as the GGDEF domain protein most frequently found to be encoded on BcsA encoding plasmids. **3C**. Phylogenetic tree of *E. coli* EAL domains of EAL domain proteins found adjacent to BcsA2-3 cellulose biosynthesis operons in the context of EAL domains from chromosomally encoded proteins. EAL domain proteins adjacent to BcsA2-3 cellulose biosynthesis operons are homologous to the chromosomally encoded core genome YliE GGDEF-EAL domain protein over the entire length of the amino acid sequence (Figure S5). Other EAL domain proteins are homologous to PdeW and Q8GGX6 accessory EAL domain protein. In addition, an EAL domain protein with a novel N-terminal signaling protein has been identified. Numbers indicate the BcsA cellulose synthase class adjacent to the EAL domain protein. Green letter; YliE GGDEF-EAL domain protein most frequently found encoded adjacent to *bcs* gene clusters; black letter, EAL domain of core genome encoded EAL domain protein of *E. coli* and *Salmonella typhimurium* (SALTY); blue letter, accessory EAL domain proteins experimentally investigated; brown letter, accessory EAL domain proteins not yet investigated. **3D**. Protter image of the chromosomally encoded CHASE9-xCache GGDEF-EAL domain protein YliE the EAL domain protein most frequently found to be encoded on BcsA encoding plasmids.

Consequently, the biological impact of two cellulose biosynthesis operon and the replacement of the core genome cellulose biosynthesis operon needs to be unravelled.

### BcsA2-BcsA4 are located on large *E. coli* plasmids

Horizontally transferred *bcsA2, bcsA3* and *bcsA4* gene clusters are located on large plasmids. Characterisation of all BcsA bearing plasmids from the genus *Escherichia* (with few from *Shigella* and *Escherichia* species other than *E. coli*) showed that the majority of plasmid belonged to the incompatibility groups p0111 and IncFIB and had a size between 39.17 kbp and 1310.597 kbp with the mean size of 143.1 ± 112.4 kbp larger than for non-BcsA encoding plasmids (mean 74.8 ± 68.0 kbp) from *E. coli* (Figure S3A). The isolation source and origin of *E. coli* strains harbouring *bcsA* bearing plasmids is divers (Figure S3B), although the majority of strains were non-pathogenic derived from environmental sources and birds collected in China and the United Kingdom. This is clearly a restricted isolation source and origin compared to all *E. coli* strains harbouring plasmids. Representative plasmids encoding *bcsA2, bcsA3* and *bcsA4* cellulose biosynthesis gene clusters are shown in Figure S2.

### GGDEF and EAL domain proteins are associated with horizontally transferred *bcs* gene clusters in *E. coli*

As a major stimulatory trigger, bacterial cellulose biosynthesis is post-translationally activated by the ubiquitous second messenger cyclic di-GMP (Ross et al. 1987, Zogaj et al. 2001, Simm et al. 2004). Diguanylate cyclase activity is conducted by the GGDEF domain, while the EAL domain has phosphodiesterase activity (Schirmer and Jenal 2009, Romling, Liang and Dow 2017). We observed that GGDEF and EAL domain protein genes commonly are located immediately adjacent to chromosomally encoded *bcs2* and *bcs3* gene clusters (Figure 3, 4 and S2), Thus we hypothesize that the co-localisation of these genes indicates connected functionality, namely that those cyclic di-GMP turnover proteins produce or hydrolise this second messenger which is required to stimulate cellulose biosynthesis. Subsequently, we investigated whether plasmids bearing *bcs* gene clusters also code for GGDEF and/or EAL domain proteins. Indeed, we identified GGDEF and EAL domain proteins encoded on *bcs* bearing plasmids (Table S2). Subsequently, we were wondering about the identity of the GGDEF and EAL domain proteins and whether they have been previously identified to regulate cellulose biosynthesis. Of note, the chromosomally encoded GGDEF and EAL domain proteins adjacent to *bcsA2* and *bcsA3* are homologous to plasmid encoded GGDEF and EAL domain proteins. However, plasmid encoded GGDEF and EAL domain proteins are more diverse.

**Figure 4.**
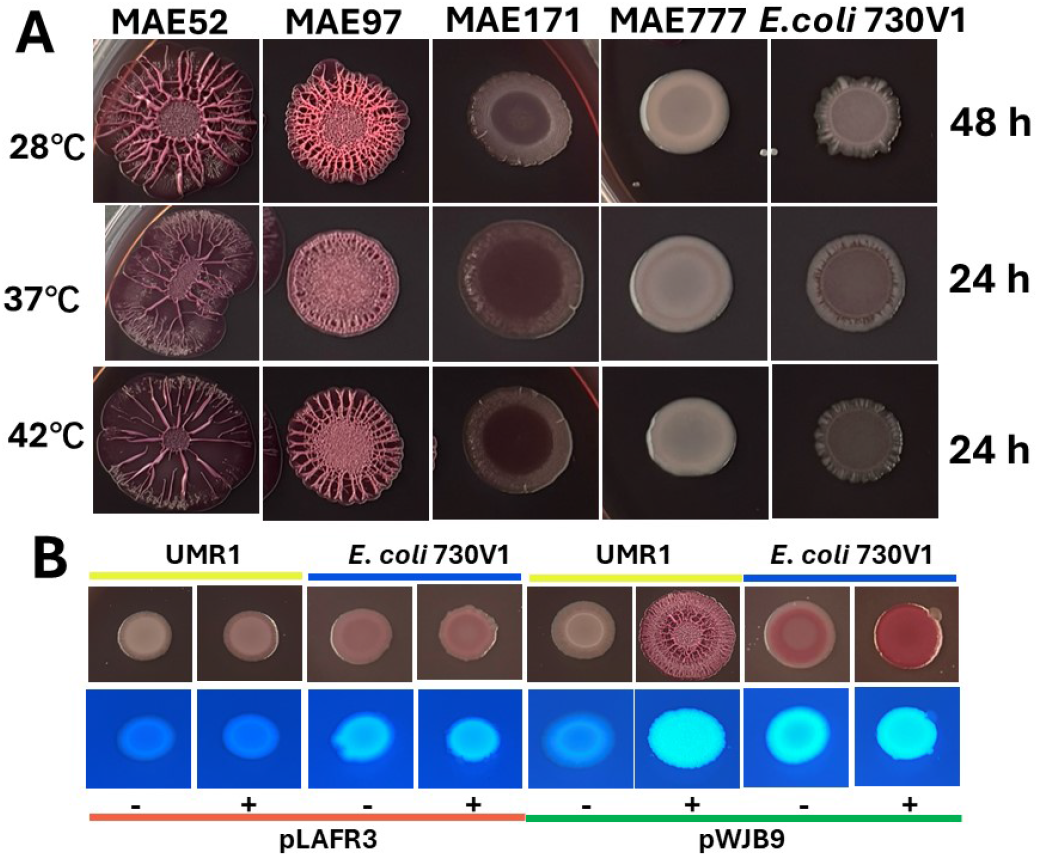
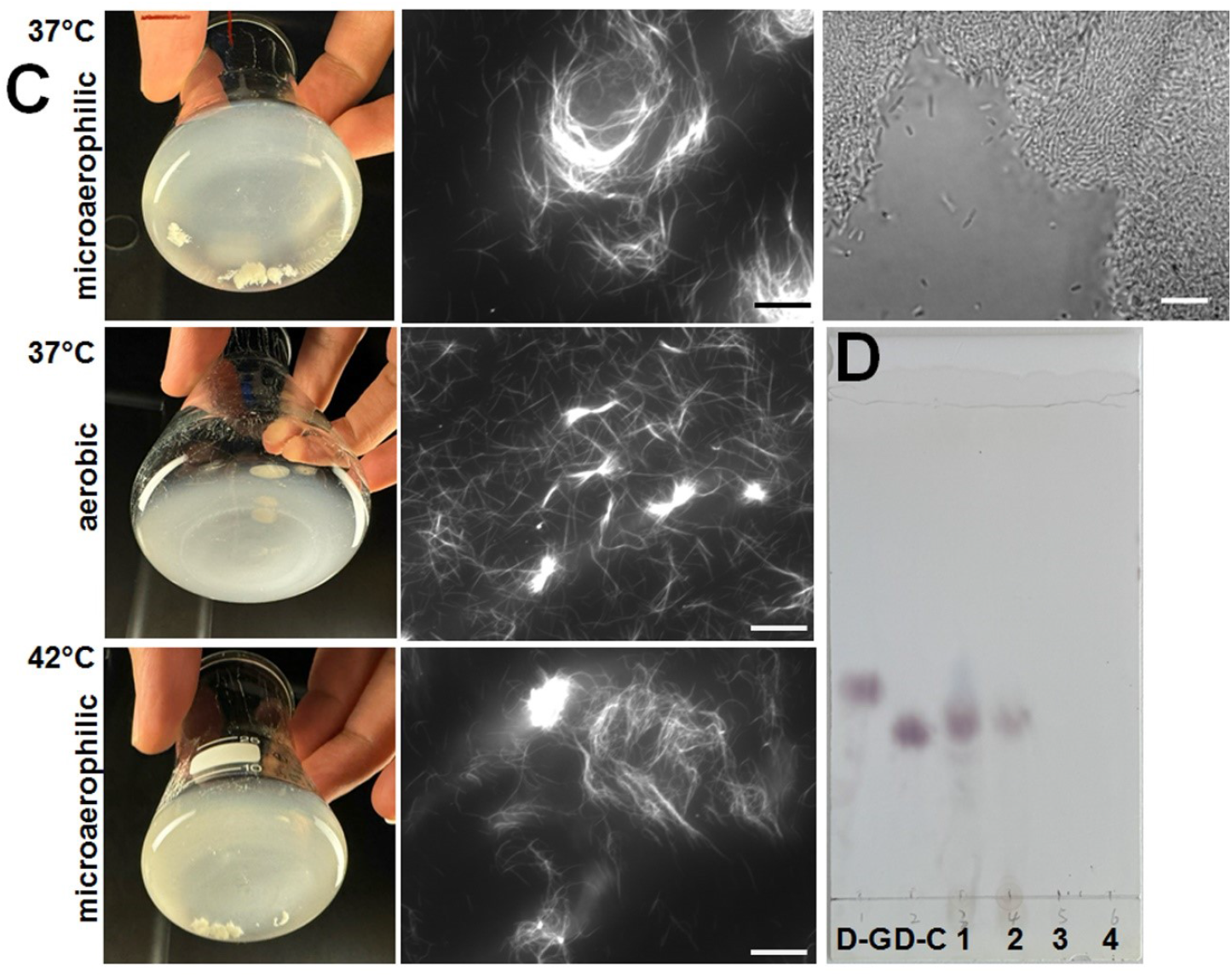
Biofilm formation and analysis of cellulose production in *E. coli* 730V1 encoding a *bcs* type 4 gene cluster on plasmid p1. **4A**. Colony morphology of *E. coli* 730V1 grown on Congo red agar plates at three relevant temperatures representing environmental, human body and meat decontamination conditions. *S. typhimurium* MAE52, a ATCC 14028 derivative (Nal^r^, P*csgD1*) and its curli (MAE97), cellulose (MAE171) and cellulose/curli negative (MAE777) mutant served as reference for expression of extracellular matrix components. **4B**. Colony morphology of *E. coli* 730V1 expressing the diguanylate cyclase AdrA from the pLAFR3 plasmid under the control of the p*BAD* promoter (pWJB9) induced by L-arabinose. *S. typhimurium* UMR1 containing pLAFR3 and pWJB9 served as reference. Cells were grown on a Congo red agar plate at 28 °C for 48 h. **4C**. Cell aggregation and Calcofluor white binding of *E. coli* 730V1 grown in M9 minimal medium with 0.4% glucose as carbon source at indicated oxygen pressure and temperature for 24 h. **4D**. Product analysis after cellulase digestion of KOH extracted exopolysaccharide from *E. coli* 730V1 grown in M9 minimal medium with 0.4% glucose as carbon source. Standard, D-G, D-glucose; D-C, cellobiose; 1, microcrystalline cellulose (Sigma-Aldrich) digested with *Trichoderma viride* cellulase; 2, KOH-extracted exopolysaccharide from *E. coli* 730V1 digested with *Trichoderma viride* cellulase; 3, *Trichoderma viride* cellulase; 4, undigested KOH-extracted exopolysaccharide from *E. coli* 730V1.

The GGDEF protein YliF, one of the *E. coli* MG1655 core genome GGDEF proteins, has the TM-GAPSE2-TM-GGDEF domain composition. *YliF* homologs located adjacent to chromosomal *bcs2* and *bcs3* operons are most frequently found on *bcsA* bearing plasmids with the three major variant 39, 32 and 41% identical over the entire length of the protein compared to YliF of MG1655 (Figure 4 A and B, S6). Conservation of amino acid motifs predict enzymatic activity for the proteins (Figure S6). In addition, truncated versions of the protein are found on several plasmids. A second GGDEF protein located on plasmids is with 83, 35 and 35 % identity over the entire length of the proteins homologs of the Globin-CBS-GGDEF protein DosC also one of the *E. coli* MG1655 GGDEF proteins.

Among the EAL domain proteins, the SP-CHASE9-xCache-TM-HAMP-xGGDEF-EAL phosphodiesterase YliE is most frequently found on *bcsA2-4* bearing plasmids and on the chromosome in several variants (Figure 4 C and D, S6). The three major YliE clades of full length proteins are 35, 37 and 28% identical to YliE of *E. coli* MG1655 and were previously identified as plasmid located (Romling, Cao and Bai 2023). As on the *E. coli* MG1655 chromosome, *yliE* is located adjacent to *yliF* at some of the *bcs* gene clusters. Besides YliE, variants of the EAL protein PdeW also previously identified on plasmids are plasmid located. In addition, variants of UrfM (QGGX6) originally identified on transposon Tn21 and a novel GGDEF protein with an unidentified N-terminal signaling domain are found on *bcsA*-bearing plasmids.

### Also other species harbor cellulose biosynthesis gene clusters on plasmids

When assessing the occurrence of the BcsA cellulose synthase on plasmids, we noticed that also plasmids contained by species isolates other than *E. coli* encode cellulose synthases with proteins closely or distantly related to the *E. coli* core genome cellulose synthase BcsA1 (Figure S1, Table S1). Thus, cellulose biosynthesis is subject to horizontal gene transfer in a larger context. Most of the BcsA bearing plasmids were harboured by *Rhizobium* spp. and *Sino-(Para- and Neo-)rhizobium* spp. isolates, alpha-proteobacterial plant symbionts known to use cellulose production for bacterial host interactions (Napoli, Dazzo and Hubbell 1975, Robledo et al. 2012). *Ensifier* spp. *Azospirillium* spp, *Caballeronia* spp. and *Klebsiella* spp. isolates are also frequently found to harbour BcsA bearing plasmids.

### Identification of bacterial species that harbour more than one cellulose biosynthesis operon

*Komagataeibacter* spp., *Proteus* spp. and *Klebsiella* spp are well known to harbour more than one cellulose biosynthesis gene cluster (Liu et al. 2020, Saxena and Brown 1995, Romling and Galperin 2015). In fact *Komagataeibacter* spp. can harbour up to four distinct *bcs* gene clusters (Ryngajllo et al. 2019). Some of these cellulose syntheses can be highly similar indicating gene duplication rather than horizontal gene transfer. As BcsA cellulose synthase are often found on plasmids, we were wondering whether those genes can become manifested on chromosomes as second cellulose biosynthesis entities. Using the Uniprot protein database, we investigated which species (or distinct isolates of a species) harbour more than one cellulose biosynthesis. In particular’, *Klebsiella* spp. and *Pseudomonas* spp. isolates can harbour more than on cellulose synthase often with low similarity indicating horizontal transfer from a phylogenetically unrelated species (Figure S7A). Surprisingly though, the second cellulose biosynthesis gene cluster is inserted immediately adjacent to the first cellulose biosynthesis gene cluster *K. pneumoniae* isolates and can also found as a third cellulose biosynthesis gene cluster on a plasmid in *K. pneumoniae* WS29-1 (Figure S7B).

### Rdar biofilm formation of *E. coli* 730V1 harbouring a *bcs4* cellulose biosynthesis operon

Remarkably, quite a number of *E. coli* strains bear a second or an alternative *bcs* gene cluster. We identified the *Bos taurus* derived isolate *E. coli* 730V1 (Guragain et al. 2021) as a strain bearing the *bcs* type 4 cellulose biosynthesis gene cluster *bcsORABCDZ bcsEFG* on a plasmid with its chromosomal *bcs* operon deleted (Figure 2). In *E. coli* and *S. typhimurium*, the exopolysaccharide cellulose is conventionally produced as an extracellular matrix component of environmental and host-associated biofilms (Lamprokostopoulou and Romling 2022). For example, concomitantly with amyloid fimbriae, cellulose is one of the two major extracellular matrix components of the agar-grown rdar (red, <u>d</u>ry <u>a</u>nd rough) colony biofilm which impacts host- and plant-associations (Kai-Larsen et al. 2010). Rdar biofilm formation is often highly regulated in most *E. coli* and *S. typhimurium* strains being expressed at ambient temperature below 30°C, but semi-constitutive biofilm formation at human body temperature can occur (Cimdins et al. 2017, Romling et al. 1998, Cimdins-Ahne et al. 2023). We assessed rdar biofilm formation of *E. coli* 730V1 at 28°C, 37°C and 42°C on Congo red and Calcoflour white (Fluorescence Brightener 28) agar plates as this isolate derived from processed meat with a history of heat intervention for decontamination. We compared *E. coli* 730V1 with the model *S. typhimurium* MAE52 (an ATCC14028 derivative) which shows a temperature independent rdar colony biofilm expressing cellulose and amyloid curli fimbriae (Romling et al. 1998). Constructed mutants of MAE52 express a cellulose-based pdar (pink, dry and rough) colony biofilm, a curli-based bdar (brown, dry and rough) colony biofilm and a saw (smooth and white) colony without expression of cellulose and curli extracellular matrix components.

We found *E. coli* 730V1 to express a weak rdar morphotype at 37°C and 42°C, but not 28°C on Congo red agar plates (Figure 5A). In contrast, the strain strongly bound Calcofluor white (Figure 5B). Also, the strain showed strong cell aggregation in M9 minimal medium at 37°C and 42°C when grown at microaerophilic conditions while Calcofluor binding was observed also under aerobic condition without strong aggregation (Figure 5C). No substantial cell aggregation was observed in LB without salt medium at the two distinct oxygen pressures. As the rdar morphotype and Calcofluor white binding are not absolutely informative for cellulose expression, we expressed the diguanylate cyclase AdrA in *E. coli* 730V1 (Figure 5B). As a control AdrA was expressed in *S. typhimurium* UMR1. In order to enable conjugation of the *adrA* gene, we selected a nalidixic acid resistant variant of *E. coli* 730V1. Of note, this strain showed a slightly upregulated rdar morphotype (Figure S8A). Upon acquisition of the adrA bearing plasmid pLAFR3, we observed elevated Congo red binding indicative for cellulose expression in *E. coli* 730V1 as for the *S. typhimurium* UMR1 control strain. Further, we isolated crystalline cellulose with KOH (obtaining crystalline and amorphous cellulose) and the Updegraff reagent (obtaining crystalline cellulose) from *E. coli* 730V1 grown under microaerophilic conditions (Figure S8B) and subjected it to cellulase digestion. Digestion of cellulose isolated from *E. coli* 730V1 and microcrystalline cellulose with cellulase obtained a product the run at the height of the cellobiose (Figure 5D). We conclude that at least a fraction of the extracellular matrix produced in M9 minimal medium is cellulose.

## Discussion

The exopolysaccharide cellulose and structurally similar poly-glucans are widespread extracellular matrix components of microbial biofilms (Zogaj et al. 2001, Perez-Mendoza et al. 2015, Bundalovic-Torma et al. 2020). Cellulose and highly similar derivatives are produced throughout the bacterial phylogenetic tree by thermophilic bacteria, by cyanobacteria in microbial mats, but have also been refunctionalized by today’s human pathogens to trade acute virulence and cytotoxicity within immune cells versus environmental survival (Romling and Galperin 2015). Cellulose and derivatives are thus multipurpose macromolecules that can protect against environmental stresses such as UV radiation and chlorine exposure (Solano et al. 2002, Williams and Cannon 1989). As an anti-virulence factor, cellulose biosynthesis prevents adhesion and invasion of bacteria into epithelial cell line and restricts the proliferation of *Salmonella* in professional macrophages (Pontes et al. 2015, Lamprokostopoulou and Romling 2022). Consequently, enhanced invasiveness and acute virulence of pathogens are associated with diminished or deleted cellulose production (Singletary et al. 2016, Nhu et al. 2024). In which ecological context a second/alternative cellulose biosynthesis operon can provide an advantage for *E. coli* remains to be investigated.

In bacteria cellulose exists not only as crystalline macrofibrills of bundles of linear parallel unmodified 1,4-beta-D-glucan chains. In several instances, the glucose subunits have been covalently modified such as in the case of class II cellulose biosynthesis operons, where glucose upon incorporation into the nascent glucan chain is covalently modified by the addition of phosphoethanolamine to the C6 carbon (Thongsomboon et al. 2018, Sun et al. 2018). The multifunctional alkaline phosphatase superfamily member BcsG not only catalyzes this transfer reaction from the most abundant phospholipid phosphoethanoleamine to glucose by its periplasmic catalytic domain, but also stabilizes the BcsA cellulose synthase via its N-terminal transmembrane part (Sun et al. 2018, Thongsomboon et al. 2018). The presence of solely the *bcsABC* gene cluster as a second copy suggests that those *E. coli* strains can produce unmodified crystalline cellulose and do not require stabilization by BcsG.

Despite the obvious impact of cellulose biosynthesis in the protection against environmental stresses, there is an open question why bacterial species (besides achieving a higher production which might in any case be limited by physiological and metabolic constraints) possess more than one cellulose biosynthesis gene cluster and whether and, if so, how these clusters are horizontally transferred or duplicated. In the fruit degrading genus *Komagataeibacter* spp with *Komagataeibacter xylinus*, model organisms for cellulose biosynthesis and the only bacterial genus that produces economically relevant amounts of cellulose has been found to encode up to four cellulose biosynthesis operons (Ryngajllo et al. 2019). Those cellulose biosynthesis operons are usually all of the same class I, but associated with different accessory gene products. However, one cellulose biosynthesis operon seems to be dominant in the production of cellulose (Saxena and Brown 1995).

Considering that the two homologous cellulose biosynthesis gene clusters in *Klebsiella* and *Proteus* spp. show only low similarity and even belong to different classes, one can hypothesize that one of these cellulose biosynthesis operons has been horizontally transferred and subsequently integrated into the chromosomes.The impact of multiple cellulose biosynthesis operons in other bacteria is, however, not that obvious. For example, although *Klebsiella pneumoniae* possesses two co-localized cellulose biosynthesis operons of different classes *K. pneumoniae* conventionally displays a mucoid colony type under laboratory conditions. Only rarely, cellulose expressing pdar colony morphology biofilms have been observed (Zogaj et al. 2001), in particular, recently causing outbreaks in hospitals in China due to their high biofilm formation capacity (Xiang et al. 2024). Of note, although possessing only one GGDEF diguanylate cyclase with cyclic di-GMP required for post-translational activation of cellulose biosynthesis, *Proteus* spp. possess two cellulose biosynthesis operons. However, cellulose biosynthesis in Proteus spp. has not been observed, even upon overexpression of a functional diguanylate cyclase (Liu et al. 2020).

The situation seems to be somewhat different in *E. coli*. Already present in the common ancestor of *Salmonella* and *E. coli*, the chromosomal location of the cellulose biosynthesis operon is highly conserved. Although diverse second cellulose biosynthesis operons have been observed on plasmids in this work, a second cellulose biosynthesis operon has not been widely established on the chromosome or on plasmids. Thus, although potentially providing a temporary and spatial niche advantage, the acquisition of a second operon does not seem to be evolutionary favorable in *E. coli* on the population level.

In contrast, the poly-N-acetyglucoseamine (PNAG) biosynthesis operon, synthesizing a second major exopolysaccharide of *E. coli* relevant for biofilm formation originally characterized in *E. coli* K-12 (Itoh et al. 2008), is not present in all *E. coli* strains and has a variable chromosomal position in the different strains (Cimdins et al. 2017). Thus, we hypothesized that the PNAG biosynthesis operon can be equally subject to horizontal gene transfer. Indeed, Blast search using PgaC showed that that the *PgaABCD* operon can be found on *E. coli* mobile elements such as plasmids (Figure S9;Table S3). In addition, *pgaABCD* operons highly similar to the chromosomally encoded PNAG operon are found on phage-like mobile elements suggesting their recent mobilization from the chromosome in different *E. coli* strains. In conclusion, even seemingly conserved biofilm components are more mobile than previously thought, but are not necessarily established population-wide in the chromosomal context.

The major regulatory mechanism to promote enhanced cellulose biosynthesis in *E. coli* is the post-translational relieve of inhibition of catalytic activity by the binding of the second messenger cyclic di-GMP to the C-terminal PilZ domain of bacterial cellulose synthases. Equally, PNAG production requires cyclic di-GMP. Consequently, we found that cyclic di-GMP turnover proteins, GGDEF diguanylate cyclases or phosphodiesterases can be encoded by plasmids encoding a cellulose biosynthesis operon (Figure S2). Thereby, the homologs of the diguanylate cyclases YliF and DosC, most frequently associated with the *bcs* operons on plasmids, have not been previously demonstrated to stimulate cellulose biosynthesis in the chromosomal context. Interestingly, though, those diguanylate cyclases have a low similarity over the entire length of the protein suggesting different regulatory and catalytic features. As Blast reach indicates that these homologous proteins, equally as the BcsA2 and BcsA3 cellulose synthases, seem to be restricted to the *Escherichia* genus, one hypothesis is that mobilization on plasmids and horizontal transfer promotes sequence diversification within the genus. Alternatively, conservation of domain structure associated with diversification has occurred in distantly related (unidentified) species and genes were subsequently mobilized.

## Conclusions

Besides the loss of cellulose biosynthesis associated with enhanced invasiveness and virulence of *E. coli* isolates, a second cellulose biosynthesis operon can also be acquired with occasional loss of the core genome encoded cellulose biosynthesis operon. While such a cellulose biosynthesis operon can be functional as shown upon post-translational activation by cyclic di-GMP in the meat-derived strain

*E. coli* 730V1, the evolutionary drivers of acquisition and, potentially maintenance of a second (and/or different cellulose biosynthesis operon have not yet been unravelled.

## Acknowledgement

We thank Dr. Joseph M Bosilevac and Dr. Manita Guragain from the Agricultural Research Service, U.S. Department of Agriculture, Meat Safety and Quality Research Unit for generously providing the *E. coli* 730V1 isolate. Part of this work has been presented at the IUMS 2024 Congress, the condrence of the International Union of Microbiological Sciences (IUMS) in Florence, Italy, October 23-25, 2024.

## Materials and Methods

### Bacterial strains and growth conditions

The following strains were used in this work. High temperature tolerant *E. coli* 730V1 isolated from beef (Guragain et al. 2021) which harbours a Bcs type 4 cellulose biosynthesis gene cluster on its large plasmid. Reference strains were *S. typhimurium* UMR1 (ATCC 14028 Nal^r^ rdar_28_), its temperature independent variant MAE52 (P*csgD1*) (Romling et al. 1998) and the respective only cellulose (MAE97), only curli (MAE171) and curli and cellulose negative (MAE777) mutants. Tetracycline was added at 20 µg/ml, kanamycin at 50 µg/ml, nalidixic acid at 50 µg/ml and L-arabinose at 0.1 % (w/v), if required.

### Selection of an *E. coli* 730V1 nalidixic acid resistant variant

*E. coli* 730V1 cells were incubated overnight in LB medium on 37°C and subsequently centrifuged at 7000 rpm for 1 min. 200 µl of an OD_600_=10 adjusted cell suspension was spread on a LB agar selection plate containing 50 *μ*g/mL nalidixic acid. After incubation for three days, two of the few colonies that grew on the selection plate were twice streaked to single colonies and subsequently stored at -80 °C as a glycerol stock. The resulting nalidixic acid resistant strain was subsequently investigated for rdar morphotype expression.

### Plasmid conjugation by triparental mating

Five hundred µl of overnight cultures of the donor strain *E. coli* containing pWJB9 (pLAFR3 containing the diguanylate cyclase AdrA under the pBAD promoter (Simm et al. 2004)) or the pLAFR3 vector control, *E. coli* containing the helper plasmid pRK2013 and the recipient strain *E. coli* 730V1 Nal^r^ were mixed and subsequently centrifuged at 13000 rpm for 1 min. The pellet was resuspended and 50 µl spotted onto an LB plate without antibiotics for overnight incubation at 37°C. Cells were subsequently resuspended in 10 mM MgSO_4_, washed once and spread onto an LB agar plate containing nalidixic acid (50 *μ*g/mL) and tetracycline (10 *μ*g/mL). The plate was incubated for three days at 37°C to recover plasmid-containing strain *E. coli* 730V1 Nal^r^ colonies. At least two of them were streaked for single colonies on the LB plate containing the antibiotics at least twice for purification.

### Bioinformatic analyses

Complete plasmids of microbes (NCBI microbial database and the curated NCBI database at https://ccb-microbe.cs.uni-saarland.de/plsdb2025) and the NCBI database of complete and draft genomes were searched for homologs to the core genome cellulose synthase BcsA1 (P37653) of *E. coli* MG1655 using default parameters. All non-redundant homologs on plasmids equally as distantly related homologs on *E. coli* chromosomes with a query coverage of > 47% and 40% were retrieved (Altschul et al. 1990). Truncated proteins were removed from the analysis. Proteins were aligned using ClustalX 2.1 with the alignment to be manually curated in GeneDoc (Higgins and Sharp 1988). Identity and similarity values for proteins were calculated using the .fasta version of the alignments with ‘Ident and Sim’ with similar amino acid classified according to GAVLI, FYW, CM, ST, KRH, DENQ, P at https://www.bioinformatics.org/sms2/ident_sim.html (Stothard 2000). For phylogenetic analysis, protein sequences were trimmed to contain the homologous regions common to all proteins. Protein alignments were displayed by ESPript 3.0 (Robert and Gouet 2014). Maximum-Likelyhood pylogenetic trees were constructed with MEGA 11.0 (Tamura, Stecher and Kumar 2021). The BcsA cellulose synthases were subsequently categorized according to the degree of homology in three additional classes, BcsA2, BcsA3 and BcsA4 and, if required, their origin as cellulose synthases of *E. coli* was verified by the phylogenetic assessment of conserved proteins informative on the species level such as GyrA. BcsA2 (WP_074014783.1) from strain *E. coli* S56 on plasmid pA, BcsA3 (HDP8224287.1) from strain *E. coli* ST-2070 (identical to the one from *E. coli* 21-X O621GT02-EC) and BcsA4 (WP_004186066.1) from strain *E. coli* 730V1 on plasmid p1 were chosen as representative proteins.

Reference BcsA cellulose synthases include class I-IV cellulose synthases (Romling and Galperin 2015), cellulose synthases most closely related to E. coli BcsA2-BcsA4 cellulose synthases and additional more distantly related cellulose synthases from the database as published previously (Cao et al. 2022). A similar strategy to retrieve homologous proteins was applied for all alternative cellulose biosynthesis proteins identified in E. coli genomes besides the core cellulose biosynthesis operon.

The gene neighborhood of the BcsA cellulose synthase genes was assessed manually for each non-redundantly retrieved BcsA cellulose synthase by evaluating the annotated plasmids. After identification of the cellulose biosynthesis gene cluster, the cluster +/-four genes up- and down-stream was downloaded. Cellulose biosynthesis gene clusters were comparatively visualized with Easyfig (Sullivan, Petty and Beatson 2011) with the most conserved BcsA cellulose synthase as reference.

Proteins (including cellulose biosynthesis operon gene products besides the BcsA cellulose synthase and GGDEF/EAL domain proteins) were aligned with ClustalX 2.1 and, if requiredto restrict the analysis to a single domain as in the case of GGDEF and EAL domain protiens, manually curated in Genedoc (Higgins and Sharp 1988). Multiple alignments with reference proteins were displayed with ESPript (Robert and Gouet 2014). A Maximum Likelihood phylogenetic tree was constructed in MEGA 7.0 or MEGA 11.0 using 1000 bootstrap iterations (Tamura et al. 2021).

Transmembrane helices were evaluated with TMHMM (Moller, Croning and Apweiler 2001) and TOPCONS (Tsirigos et al. 2015). TM helices and domain borders of BcsA cellulose synthases were evaluated against the crystal structure of the *Cereibacter sphaeroides* cellulose synthase (PDB: 4HG6). Conserved signature amino acids for the cellulose synthase were taken from the literature (Romling 2002, Oehme et al. 2019).

Domain structure were displayed with Illustrator for Biological Sequences IBS2.0 (Xie et al. 2022). Protter (Omasits et al. 2014) was used to visualize the amino acid sequences within the domain structures.

FimH types, serotypes and origin of replication were identified using servers provided by the Center of Genomic Epidemiology, FimTypes, SerotypeFinder and PlasmidFinder 2.1, respectively (Carattoli and Hasman 2020). All BcsA bearing plasmids were searched and retrieved from the PLSDB database at https://ccb-microbe.cs.uni-saarland.de/plsdb2025/ (Galata et al. 2019). Representative plasmids were displayed by Proksee (Grant et al. 2023) reannotated with Prokka (Seemann 2014).

### Box plot calculation with R

Mean and median plasmid size was calculated and the box plot with borders indicating the first and third quadrille drawn using R studio version 4.4.

### Assessment of cell aggregation, cellulose expression and colony biofilm formation

Bacteria were recovered from -80°C stocks by streaking onto a LB agar plate. After overnight growth, colonies were resuspended in LB without salt medium and adjusted to an OD=5. Subsequently, 5 µl of cell suspension was spotted on LB without salt agar medium supplemented with Congo red (CR) (CR 40 µg/ml and Coomassie Brilliant Blue 20 µg/ml) or Calcofluor white (fluorescence brightener 28, 50 µg/ml) and allowed to dry (Cimdins et al. 2017). Colony morphology was assessed visually or by observing the cells exposed to a 366 nm UV light source after 24, 48 and 72 h of growth at 28°C, 37°C or 42°C and documented by photographing. To induce the production of the AdrA diguanylate cyclase under the control of the pBAD promoter, 0.1% L-arabinose was added to Congo red and Calcofluor white agar plates (Simm et al. 2004).

Biofilm formation and cell aggregation in liquid culture was assessed after either aerobic or microaerophilic growth in LB without salt and M9 medium supplemented with 0.4% glucose (Gerstel and Romling 2001). Culture flasks were photographed at the end of the incubation period and bacterial aggregates were subsequently observed under a fluorescence microscope (Nikon eclipse 80i) using excitation wave length of 340-380 nm and observing emission at wavelength of 435-485 nm after staining with fluorescent brightener 28.

### Extraction of crystalline cellulose by the Updegraff reagent

Crystalline cellulose has been extracted by the Updegraff reagent (58% acetic acid, 19% nitric acid (Updegraff 1969) at 95°C for 30 min and the pellet subsequently washed with double distilled H_2_O.

### Extraction of cellulose by 1% KOH

Cells grown in M9 minimum medium with 0.4% glucose under microaerophilic conditions were harvested after 24 h incubation, resuspended in 1% potassium hydroxide and incubated for 1 h at 95 °C under strong shaking. The sample was centrifuged at 13,300 rpm for 10 min and the pellet washed three times with water.

### Analysis of cellulase-digested products by thin layer chromatography

KOH-isolated cellulose from *E. coli* 730V1 was digested with 0.1 mg/mL cellulase (from *Trichoderma viride*, Sigma) in 0.05 M sodium citrate pH 4.6 overnight. The reaction mixture was dried, dissolved in water and centrifuged at 13,000 g for 15 min. 2 *μ*L of the sample was applied on a 5 x 20 cm silica-based thin layer chromatography plate (Spelco, TLC plates, silica gel 60 F_254_). 2 µl of a glucose and cellobiose solution (5 mg/mL) was used as standard. The solvent system consisted of n-butanol, acetic acid and water (2:1:1), respectively. After drying, the plate was emersed in the dipping solution orcinol (0.2 g/100 mL) in sulfuric acid (20 g/100 mL) and heated for 10 min at 100 °C.

## Tables

**Table S1**

List of reference plasmids from Blast search used for analysis with information on description, scientific name, max score, total score, query cover e-value, percent identity, actual length of plasmid, accession, identical proteins.

**Table S2**

Description, strain, plasmid, Fimtype, O type H type, Country, infected source number of all GGDEF domais found, number of all EAL domains found, GGDEF domain protein id, EAL domain protein id.

**Table S3**

List of *pgaABCD* encoding mobile plasmidic elements in *Escherichia coli* identified by Blast search with plasmid accession number, number of hits, length [aa], ORF start, ORF stop, e-value, bitscore, identity to query, query coverage, plasmid topology, submission date, graphical origin, plasmid origin of replication (PlasmidFinder), plasmid origin of replication (pMLST) and plasmid size.

## References

Ahmad, I., S. F. Rouf, L. Sun, A. Cimdins, S. Shafeeq, S. Le Guyon, M. Schottkowski, M. Rhen & U. Romling (2016) BcsZ inhibits biofilm phenotypes and promotes virulence by blocking cellulose production in Salmonella enterica serovar Typhimurium. Microb Cell Fact, 15, 177.

Altschul, S. F., W. Gish, W. Miller, E. W. Myers & D. J. Lipman (1990) Basic local alignment search tool. J Mol Biol, 215, 403–10.

Bundalovic-Torma, C., G. B. Whitfield, L. S. Marmont, P. L. Howell & J. Parkinson (2020) A systematic pipeline for classifying bacterial operons reveals the evolutionary landscape of biofilm machineries. PLoS Comput Biol, 16, e1007721.

Cao, L. Y., Y. F. Yang, X. Zhang, Y. H. Chen, J. W. Yao, X. Wang, J. Xia, C. G. Liu, S. H. Yang, U. Romling & F. W. Bai (2022) Deciphering Molecular Mechanism Underlying Self-Flocculation of Zymomonas mobilis for Robust Production. Appl Environ Microbiol, 88, e0239821.

Carattoli, A. & H. Hasman (2020) PlasmidFinder and In Silico pMLST: Identification and Typing of Plasmid Replicons in Whole-Genome Sequencing (WGS). Methods Mol Biol, 2075, 285–294.

Chang, S. C., M. R. Kao, R. K. Saldivar, S. M. Diaz-Moreno, X. Xing, V. Furlanetto, J. Yayo, C. Divne, F. Vilaplana, D. W. Abbott & Y. S. Y. Hsieh (2023) The Gram-positive bacterium Romboutsia ilealis harbors a polysaccharide synthase that can produce (1,3;1,4)-beta-D-glucans. Nat Commun, 14, 4526.

Chanin, R. B., K. P. Nickerson, A. Llanos-Chea, J. R. Sistrunk, D. A. Rasko, D. K. V. Kumar, J. de la Parra, J. R. Auclair, J. Ding, K. Li, S. K. Dogiparthi, B. J. D. Kusber & C. S. Faherty (2019) Shigella flexneri Adherence Factor Expression in In Vivo-Like Conditions. mSphere, 4.

Cimdins-Ahne, A., A. O. Naemi, F. Li, R. Simm & U. Romling (2023) Characterisation of Variants of Cyclic di-GMP Turnover Proteins Associated with Semi-Constitutive rdar Morphotype Expression in Commensal and Uropathogenic Escherichia coli Strains. Microorganisms, 11.

Cimdins, A., R. Simm, F. Li, P. Luthje, K. Thorell, A. Sjoling, A. Brauner & U. Romling (2017) Alterations of c-di-GMP turnover proteins modulate semi-constitutive rdar biofilm formation in commensal and uropathogenic Escherichia coli. Microbiologyopen, 6.

Fang, X., I. Ahmad, A. Blanka, M. Schottkowski, A. Cimdins, M. Y. Galperin, U. Romling & M. Gomelsky (2014) GIL, a new c-di-GMP-binding protein domain involved in regulation of cellulose synthesis in enterobacteria. Mol Microbiol, 93, 439–52.

Fugelstad, J., J. Bouzenzana, S. Djerbi, G. Guerriero, I. Ezcurra, T. T. Teeri, L. Arvestad & V. Bulone (2009) Identification of the cellulose synthase genes from the Oomycete Saprolegnia monoica and effect of cellulose synthesis inhibitors on gene expression and enzyme activity. Fungal Genet Biol, 46, 759–67.

Galata, V., T. Fehlmann, C. Backes & A. Keller (2019) PLSDB: a resource of complete bacterial plasmids. Nucleic Acids Res, 47, D195–D202.

Gerstel, U. & U. Romling (2001) Oxygen tension and nutrient starvation are major signals that regulate agfD promoter activity and expression of the multicellular morphotype in Salmonella typhimurium. Environ Microbiol, 3, 638–48.

Grant, J. R., E. Enns, E. Marinier, A. Mandal, E. K. Herman, C. Y. Chen, M. Graham, G. Van Domselaar & P. Stothard (2023) Proksee: in-depth characterization and visualization of bacterial genomes. Nucleic Acids Res, 51, W484–W492.

Guragain, M., D. M. Brichta-Harhay, J. L. Bono & J. M. Bosilevac (2021) Locus of Heat Resistance (LHR) in Meat-Borne Escherichia coli: Screening and Genetic Characterization. Appl Environ Microbiol, 87.

Higgins, D. G. & P. M. Sharp (1988) CLUSTAL: a package for performing multiple sequence alignment on a microcomputer. Gene, 73, 237–44.

Itoh, Y., J. D. Rice, C. Goller, A. Pannuri, J. Taylor, J. Meisner, T. J. Beveridge, J. F. Preston, 3rd & T. Romeo (2008) Roles of pgaABCD genes in synthesis, modification, and export of the Escherichia coli biofilm adhesin poly-beta-1,6-N-acetyl-D-glucosamine. J Bacteriol, 190, 3670–80.

Jeon, Y. J., Z. Xun, P. Su & P. L. Rogers (2012) Genome-wide transcriptomic analysis of a flocculent strain of Zymomonas mobilis. Appl Microbiol Biotechnol, 93, 2513–8.

Kai-Larsen, Y., P. Luthje, M. Chromek, V. Peters, X. Wang, A. Holm, L. Kadas, K. O. Hedlund, J. Johansson, M. R. Chapman, S. H. Jacobson, U. Romling, B. Agerberth & A. Brauner (2010) Uropathogenic Escherichia coli modulates immune responses and its curli fimbriae interact with the antimicrobial peptide LL-37. PLoS Pathog, 6, e1001010.

Kamal, S. M., A. Cimdins-Ahne, C. Lee, F. Li, A. J. Martin-Rodriguez, Z. Seferbekova, R. Afasizhev, H. T. Wami, P. Katikaridis, L. Meins, H. Lunsdorf, U. Dobrindt, A. Mogk & U. Romling (2021) A recently isolated human commensal Escherichia coli ST10 clone member mediates enhanced thermotolerance and tetrathionate respiration on a P1 phage-derived IncY plasmid. Mol Microbiol, 115, 255–271.

Klemm, D., B. Heublein, H. P. Fink & A. Bohn (2005) Cellulose: fascinating biopolymer and sustainable raw material. Angew Chem Int Ed Engl, 44, 3358–93.

Lamprokostopoulou, A. & U. Romling (2022) Yin and Yang of Biofilm Formation and Cyclic di-GMP Signaling of the Gastrointestinal Pathogen Salmonella enterica Serovar Typhimurium. J Innate Immun, 14, 275–292.

Liu, Y., C. Lee, F. Li, J. Trcek, H. Bahre, R. T. Guo, C. C. Chen, A. Chernobrovkin, R. Zubarev & U. Romling (2020) A Cyclic di-GMP Network Is Present in Gram-Positive Streptococcus and Gram-Negative Proteus Species. ACS Infect Dis, 6, 2672–2687.

Menendez, E., M. Robledo, J. I. Jimenez-Zurdo, E. Velazquez, R. Rivas, J. D. Murray & P. F. Mateos (2019) Legumes display common and host-specific responses to the rhizobial cellulase CelC2 during primary symbiotic infection. Sci Rep, 9, 13907.

Moller, S., M. D. Croning & R. Apweiler (2001) Evaluation of methods for the prediction of membrane spanning regions. Bioinformatics, 17, 646–53.

Morgan, J. L., J. T. McNamara & J. Zimmer (2014) Mechanism of activation of bacterial cellulose synthase by cyclic di-GMP. Nat Struct Mol Biol, 21, 489–96.

Nakashima, K., L. Yamada, Y. Satou, J. Azuma & N. Satoh (2004) The evolutionary origin of animal cellulose synthase. Dev Genes Evol, 214, 81–8.

Napoli, C., F. Dazzo & D. Hubbell (1975) Production of cellulose microfibrils by Rhizobium. Appl Microbiol, 30, 123–31.

Nhu, N. T. K., M. A. Rahman, K. G. K. Goh, S. J. Kim, M. D. Phan, K. M. Peters, L. Alvarez-Fraga, S. J. Hancock, C. Ravi, T. J. Kidd, M. J. Sullivan, K. M. Irvine, S. A. Beatson, M. J. Sweet, A. D. Irwin, J. Vukovic, G. C. Ulett, S. Z. Hasnain & M. A. Schembri (2024) A convergent evolutionary pathway attenuating cellulose production drives enhanced virulence of some bacteria. Nat Commun, 15, 1441.

Oehme, D. P., T. Shafee, M. T. Downton, A. Bacic & M. S. Doblin (2019) Differences in protein structural regions that impact functional specificity in GT2 family beta-glucan synthases. PLoS One, 14, e0224442.

Omasits, U., C. H. Ahrens, S. Muller & B. Wollscheid (2014) Protter: interactive protein feature visualization and integration with experimental proteomic data. Bioinformatics, 30, 884–6.

Pedersen, G. B., L. Blaschek, K. E. H. Frandsen, L. C. Noack & S. Persson (2023) Cellulose synthesis in land plants. Mol Plant, 16, 206–231.

Perez-Mendoza, D., M. A. Rodriguez-Carvajal, L. Romero-Jimenez, A. Farias Gde, J. Lloret, M. T. Gallegos & J. Sanjuan (2015) Novel mixed-linkage beta-glucan activated by c-di-GMP in Sinorhizobium meliloti. Proc Natl Acad Sci U S A, 112, E757–65.

Pfeifer, E., J. A. Moura de Sousa, M. Touchon & E. P. C. Rocha (2021) Bacteria have numerous distinctive groups of phage-plasmids with conserved phage and variable plasmid gene repertoires. Nucleic Acids Res, 49, 2655–2673.

Pontes, M. H., E. J. Lee, J. Choi & E. A. Groisman (2015) Salmonella promotes virulence by repressing cellulose production. Proc Natl Acad Sci U S A, 112, 5183–8.

Robert, X. & P. Gouet (2014) Deciphering key features in protein structures with the new ENDscript server. Nucleic Acids Res, 42, W320–4.

Robledo, M., L. Rivera, J. I. Jimenez-Zurdo, R. Rivas, F. Dazzo, E. Velazquez, E. Martinez-Molina, A. M. Hirsch & P. F. Mateos (2012) Role of Rhizobium endoglucanase CelC2 in cellulose biosynthesis and biofilm formation on plant roots and abiotic surfaces. Microb Cell Fact, 11, 125.

Romling, U. (2002) Molecular biology of cellulose production in bacteria. Res Microbiol, 153, 205–12.

Romling, U., L. Y. Cao & F. W. Bai (2023) Evolution of cyclic di-GMP signalling on a short and long term time scale. Microbiology (Reading), 169.

Romling, U. & M. Y. Galperin (2015) Bacterial cellulose biosynthesis: diversity of operons, subunits, products, and functions. Trends Microbiol, 23, 545–57.

Romling, U., Z. X. Liang & J. M. Dow (2017) Progress in Understanding the Molecular Basis Underlying Functional Diversification of Cyclic Dinucleotide Turnover Proteins. J Bacteriol, 199.

Romling, U., M. Rohde, A. Olsen, S. Normark & J. Reinkoster (2000) AgfD, the checkpoint of multicellular and aggregative behaviour in Salmonella typhimurium regulates at least two independent pathways. Mol Microbiol, 36, 10–23.

Romling, U., W. D. Sierralta, K. Eriksson & S. Normark (1998) Multicellular and aggregative behaviour of Salmonella typhimurium strains is controlled by mutations in the agfD promoter. Mol Microbiol, 28, 249–64.

Ross, P., H. Weinhouse, Y. Aloni, D. Michaeli, P. Weinberger-Ohana, R. Mayer, S. Braun, E. de Vroom, G. A. van der Marel, J. H. van Boom & M. Benziman (1987) Regulation of cellulose synthesis in Acetobacter xylinum by cyclic diguanylic acid. Nature, 325, 279–81.

Ryngajllo, M., K. Kubiak, M. Jedrzejczak-Krzepkowska, P. Jacek & S. Bielecki (2019) Comparative genomics of the Komagataeibacter strains-Efficient bionanocellulose producers. Microbiologyopen, 8, e00731.

Sana, T. G., A. Notopoulou, L. Puygrenier, M. Decossas, S. Moreau, A. Carlier & P. V. Krasteva (2024) Structures and roles of BcsD and partner scaffold proteins in proteobacterial cellulose secretion. Curr Biol, 34, 106–116 e6.

Saxena, I. M. & R. M. Brown, Jr. (1995) Identification of a second cellulose synthase gene (acsAII) in Acetobacter xylinum. J Bacteriol, 177, 5276–83.

Schirmer, T. & U. Jenal (2009) Structural and mechanistic determinants of c-di-GMP signalling. Nat Rev Microbiol, 7, 724–35.

Seemann, T. (2014) Prokka: rapid prokaryotic genome annotation. Bioinformatics, 30, 2068–9.

Simm, R., M. Morr, A. Kader, M. Nimtz & U. Romling (2004) GGDEF and EAL domains inversely regulate cyclic di-GMP levels and transition from sessility to motility. Mol Microbiol, 53, 1123–34.

Singletary, L. A., J. E. Karlinsey, S. J. Libby, J. P. Mooney, K. L. Lokken, R. M. Tsolis, M. X. Byndloss, L. A. Hirao, C. A. Gaulke, R. W. Crawford, S. Dandekar, R. A. Kingsley, C. L. Msefula, R. S. Heyderman & F. C. Fang (2016) Loss of Multicellular Behavior in Epidemic African Nontyphoidal Salmonella enterica Serovar Typhimurium ST313 Strain D23580. mBio, 7, e02265.

Solano, C., B. Garcia, J. Valle, C. Berasain, J. M. Ghigo, C. Gamazo & I. Lasa (2002) Genetic analysis of Salmonella enteritidis biofilm formation: critical role of cellulose. Mol Microbiol, 43, 793–808.

Spiers, A. J., J. Bohannon, S. M. Gehrig & P. B. Rainey (2003) Biofilm formation at the air-liquid interface by the Pseudomonas fluorescens SBW25 wrinkly spreader requires an acetylated form of cellulose. Mol Microbiol, 50, 15–27.

Stothard, P. (2000) The sequence manipulation suite: JavaScript programs for analyzing and formatting protein and DNA sequences. Biotechniques, 28, 1102, 1104.

Sullivan, M. J., N. K. Petty & S. A. Beatson (2011) Easyfig: a genome comparison visualizer. Bioinformatics, 27, 1009–10.

Sun, L., P. Vella, R. Schnell, A. Polyakova, G. Bourenkov, F. Li, A. Cimdins, T. R. Schneider, Y. Lindqvist, M. Y. Galperin, G. Schneider & U. Romling (2018) Structural and Functional Characterization of the BcsG Subunit of the Cellulose Synthase in Salmonella typhimurium. J Mol Biol, 430, 3170–3189.

Tamura, K., G. Stecher & S. Kumar (2021) MEGA11: Molecular Evolutionary Genetics Analysis Version 11. Mol Biol Evol, 38, 3022–3027.

Thongsomboon, W., D. O. Serra, A. Possling, C. Hadjineophytou, R. Hengge & L. Cegelski (2018) Phosphoethanolamine cellulose: A naturally produced chemically modified cellulose. Science, 359, 334–338.

Tsirigos, K. D., C. Peters, N. Shu, L. Kall & A. Elofsson (2015) The TOPCONS web server for consensus prediction of membrane protein topology and signal peptides. Nucleic Acids Res, 43, W401–7.

Updegraff, D. M. (1969) Semimicro determination of cellulose in biological materials. Anal Biochem, 32, 420–4.

Verma, P., R. Ho, S. A. Chambers, L. Cegelski & J. Zimmer (2024) Insights into phosphoethanolamine cellulose synthesis and secretion across the Gram-negative cell envelope. Nat Commun, 15, 7798.

Williams, W. S. & R. E. Cannon (1989) Alternative Environmental Roles for Cellulose Produced by Acetobacter xylinum. Appl Environ Microbiol, 55, 2448–52.

Wong, H. C., A. L. Fear, R. D. Calhoon, G. H. Eichinger, R. Mayer, D. Amikam, M. Benziman, D. H. Gelfand, J. H. Meade, A. W. Emerick & et al. (1990) Genetic organization of the cellulose synthase operon in Acetobacter xylinum. Proc Natl Acad Sci U S A, 87, 8130–4.

Xiang, Y., K. Zhu, K. Min, Y. Zhang, J. Liu, K. Liu, Y. Han, X. Li, X. Du, X. Wang, Y. Huang, X. Li, Y. Peng, C. Yang, H. Liu, H. Liu, X. Li, H. Wang, C. Wang, Q. Wang, H. Jia, M. Yang, L. Wang, Y. Wu, Y. Cui, F. Chen, H. Yang, S. Baker, X. Xu, J. Yang, H. Song & S. Qiu (2024) Characterization of a Salmonella enterica serovar Typhimurium lineage with rough colony morphology and multidrug resistance. Nat Commun, 15, 6123.

Xie, Y., H. Li, X. Luo, H. Li, Q. Gao, L. Zhang, Y. Teng, Q. Zhao, Z. Zuo & J. Ren (2022) IBS 2.0: an upgraded illustrator for the visualization of biological sequences. Nucleic Acids Res, 50, W420–W426.

Zogaj, X., M. Nimtz, M. Rohde, W. Bokranz & U. Romling (2001) The multicellular morphotypes of Salmonella typhimurium and Escherichia coli produce cellulose as the second component of the extracellular matrix. Mol Microbiol, 39, 1452–63.

