## Supplementary data for "Diverse horizontally transferred cellulose biosynthesis gene clusters in *Escherichia coli* strains"

**Content**

**Figures	page**

Figure S1A-D	3

Figure S2	9

Figure S3	10

Figure S4	11

Figure S5	13

Figure S6A-C	19

Figure S7A,B	22

Figure S8A,B	25

Figure S9	26

**Tables**

Table S1

Table S2

Table S4

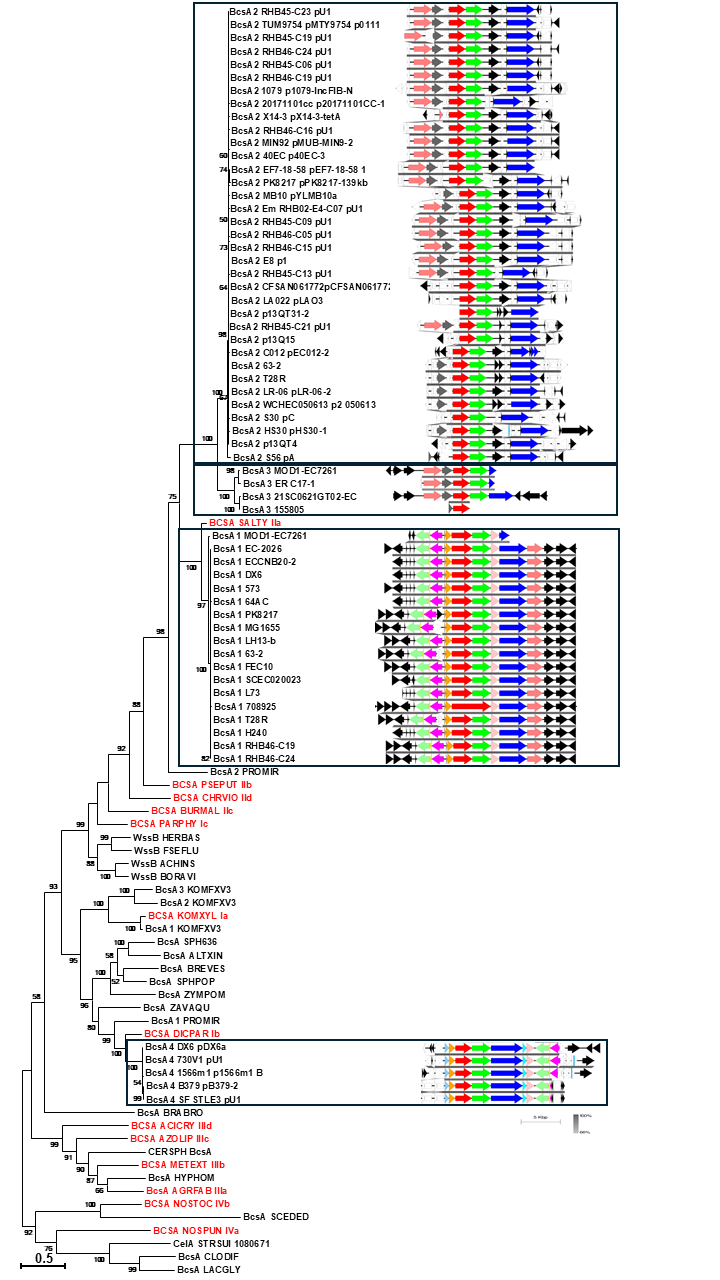

**Figure S1**

**S1A.** Extended phylogenetic tree of BcsA proteins from *E. coli* and BcsA reference proteins including the genomic context. BcsA cellulose synthases (see appendix and Figure S1A) were aligned with ClustalX and subsequently manually curated. The phylogenetic relationship has been assessed using a maximum likelihood (ML) based phylogenetic tree with 1000 bootstraps in MEGA 11.0. The nucleotide sequences and annotations of the *bcs* gene clusters and their gene neighborhoods were extracted from the genomic sequences stored at NCBI and aligned and visualized with Easyfig (Sullivan et al. 2011). Gene annotation, see Figure 1. Size bar, substitutions per site.

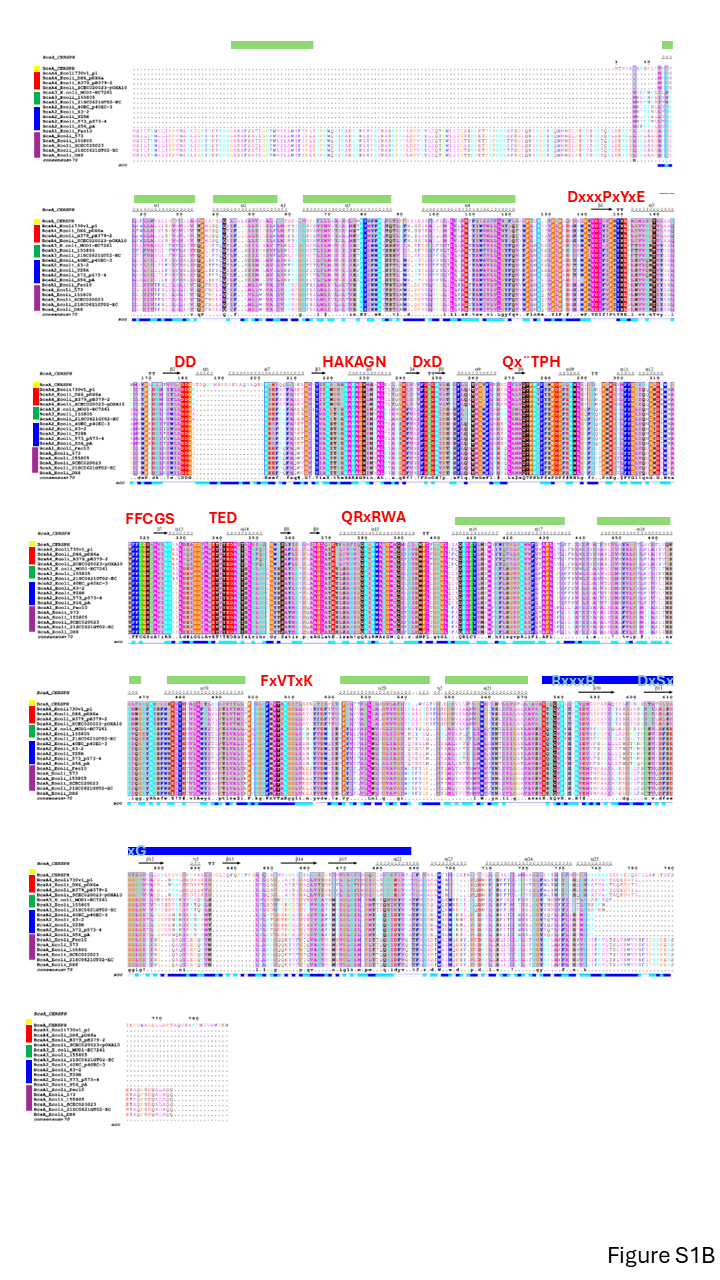

**S1B.** Alignment of representative BcsA2-4 cellulose synthases from *E. coli* with reference proteins including BcsA1 cellulose syntheases of *E. coli.* Conserved motifs and transmembrane helices characteristic for glycosyltransferases including cellulose synthases are indicated.

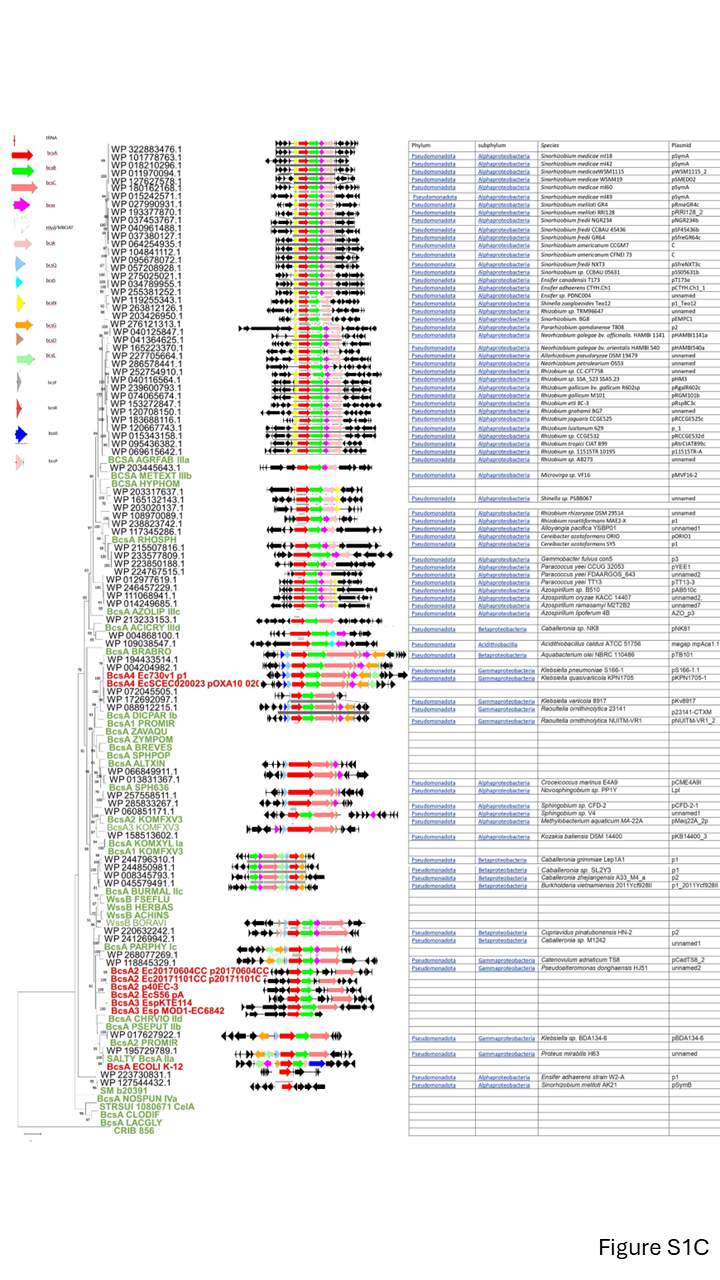

**S1C.** Phylogenetic tree of BcsA proteins of non-*E. coli* plasmids with BcsA protein from *E. coli* and BcsA reference proteins including the genomic context. BcsA cellulose synthases (see appendix and Figure S1A) were aligned with ClustalX and subsequently manually curated. The phylogenetic relationship has been assessed using a maximum likelihood (ML) based phylogenetic tree with 1000 bootstraps in MEGA 11.0. The nucleotide sequences and annotations of the *bcs* gene clusters and their gene neighborhoods were extracted from the genomic sequences stored at NCBI and aligned and visualized with Easyfig (Sullivan et al. 2011).

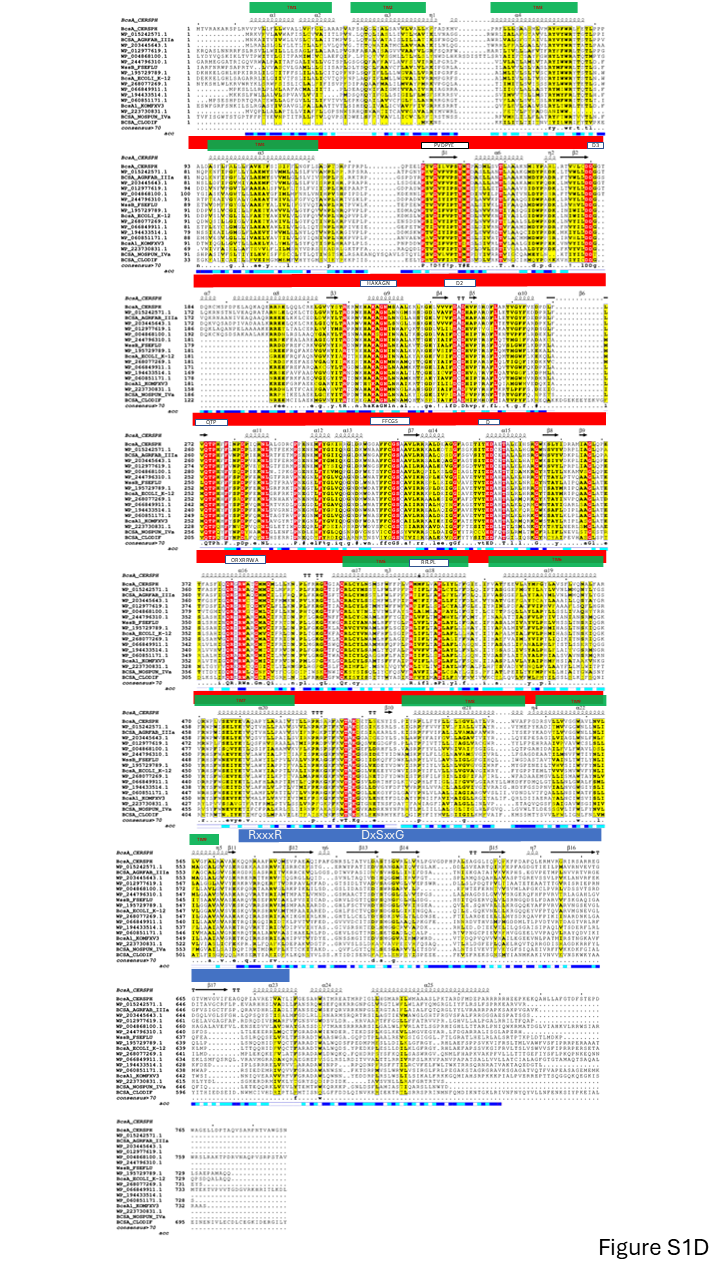

**S1D.** Alignment of cellulose synthases from non-*E. coli* plasmids with reference BcsA proteins*.* Conserved motifs and transmembrane helices characteristic for glycosyltransferases including cellulose synthases are indicated.

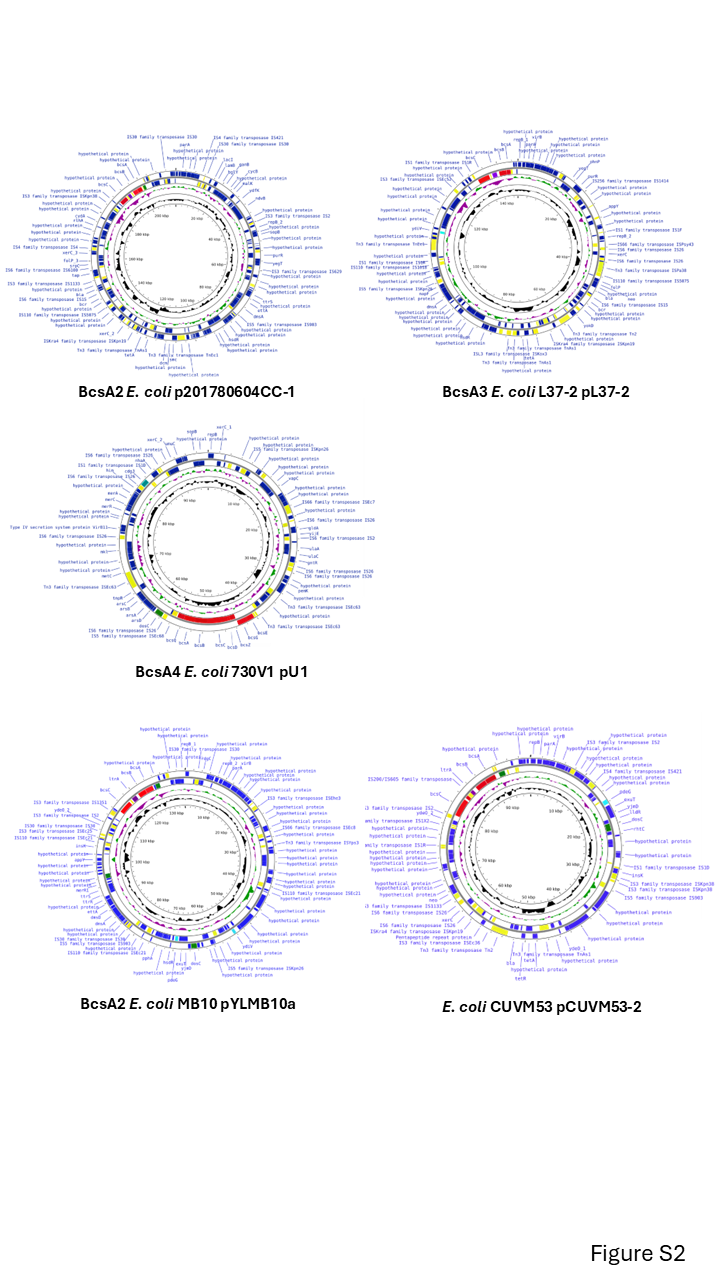

**Figure S2**

Representative plasmids of the different classes harbouring *bcsA2-4* cellulose biosynthesis gene clusters and plasmids with numerous GGDEF/EAL domain proteins (*E. coli* MB10 pYLMB10a and *E. coli* CUVM53 pCUVM53-2). In red, cellulose biosynthesis genes; green, GGDEF diguanylate cyclases; cyan, EAL phosphodisterases; yellow, transposases. Plasmids were visualized with Proksee.

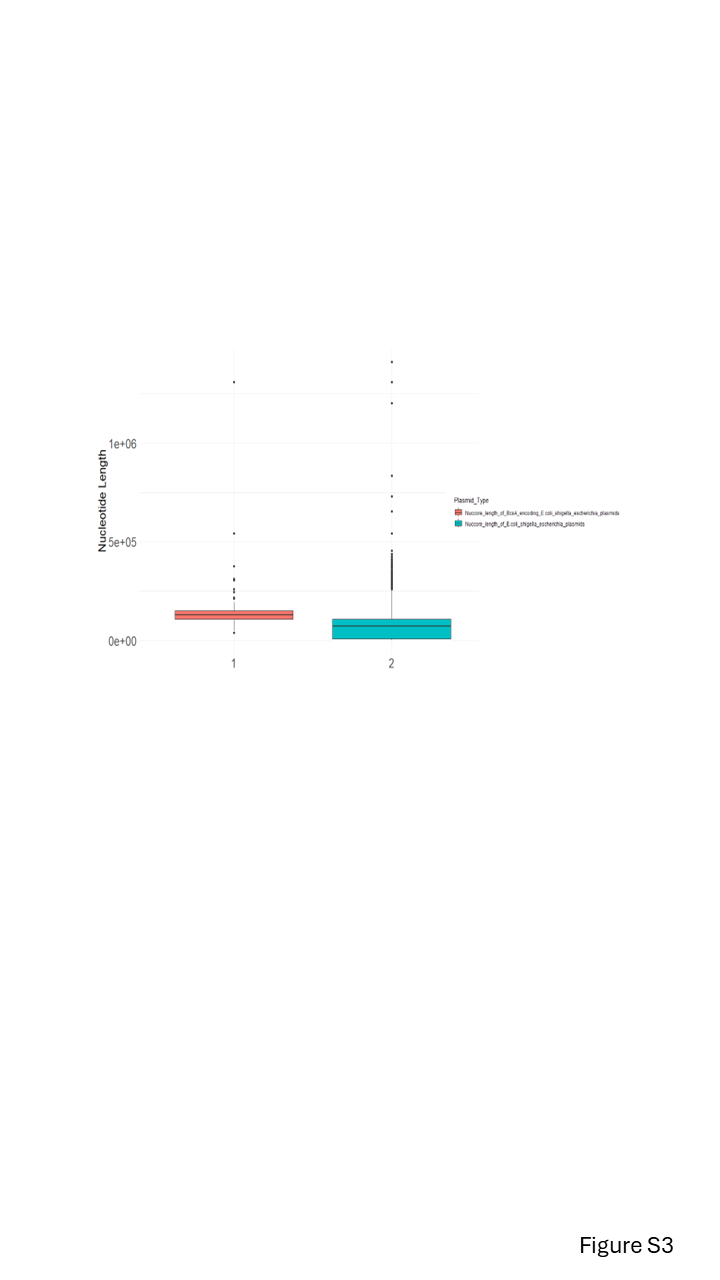

**Figure S3**

Size distribution of BcsA bearing plasmids versus all plasmids of *E. coli*, *Shigella* and *Escherichia* spp. as calculated with R studio version 4.4.

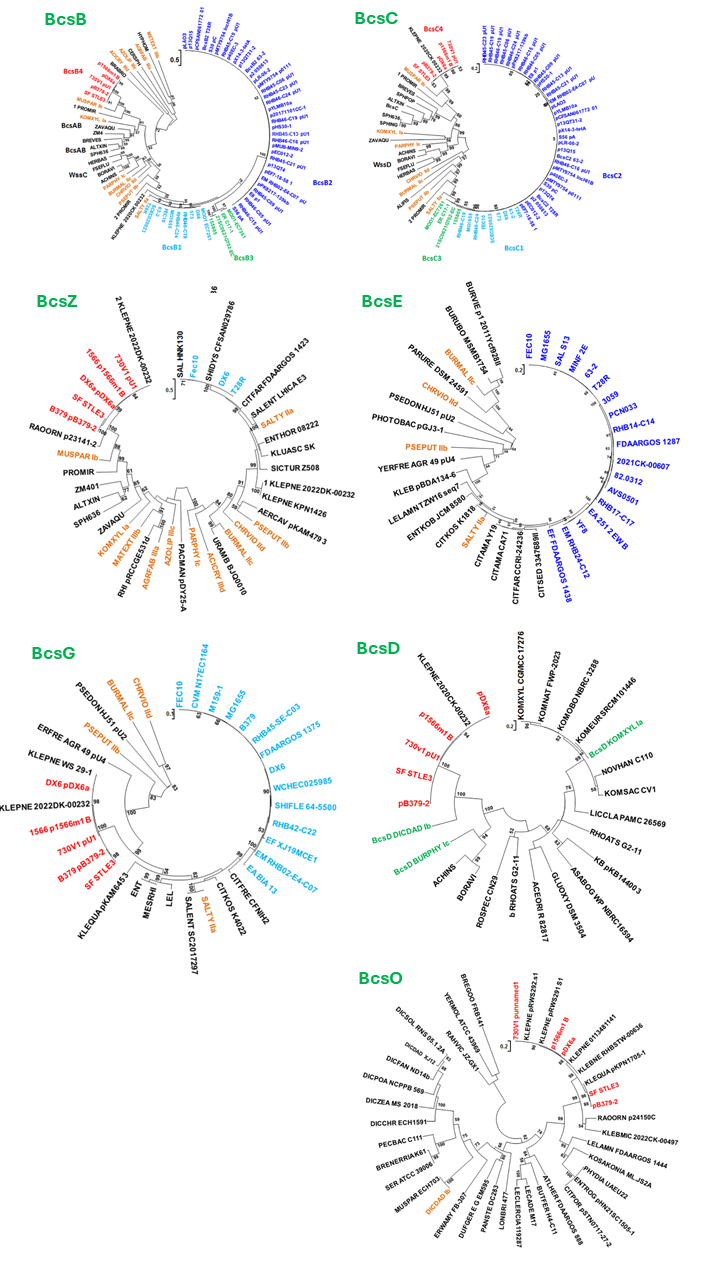

**Figure S4**

Phylogenetic trees of BcsB, BcsC, BcsE, BcsG, BcsZ, BcsD and BcsO proteins of cellulose biosynthesis operons from BcsA encoding *E. coli* plasmids and in the phylogenetic context. Proteins were retrieved from the respective BcsA2-4 cellulose biosynthesis operons and representative proteins obtained by BLAST homology search. Proteins were aligned with ClustalX 2.1 and subsequently manually curated. The phylogenetic relationship has been assessed using a maximum likelihood (ML) based phylogenetic tree with 1000 bootstraps calculated and created in MEGA 11.0. Light blue designation, protein from *E. coli* type 1 *bcs* gene cluster; dark blue, protein from *E. coli* type 2 *bcs* gene cluster; green, protein from *E. coli* type 3 *bcs* gene cluster red, protein from *E. coli* type 4 *bcs* gene cluster; brown, protein from reference *bcs* gene cluster. Size bar, substitution per site.

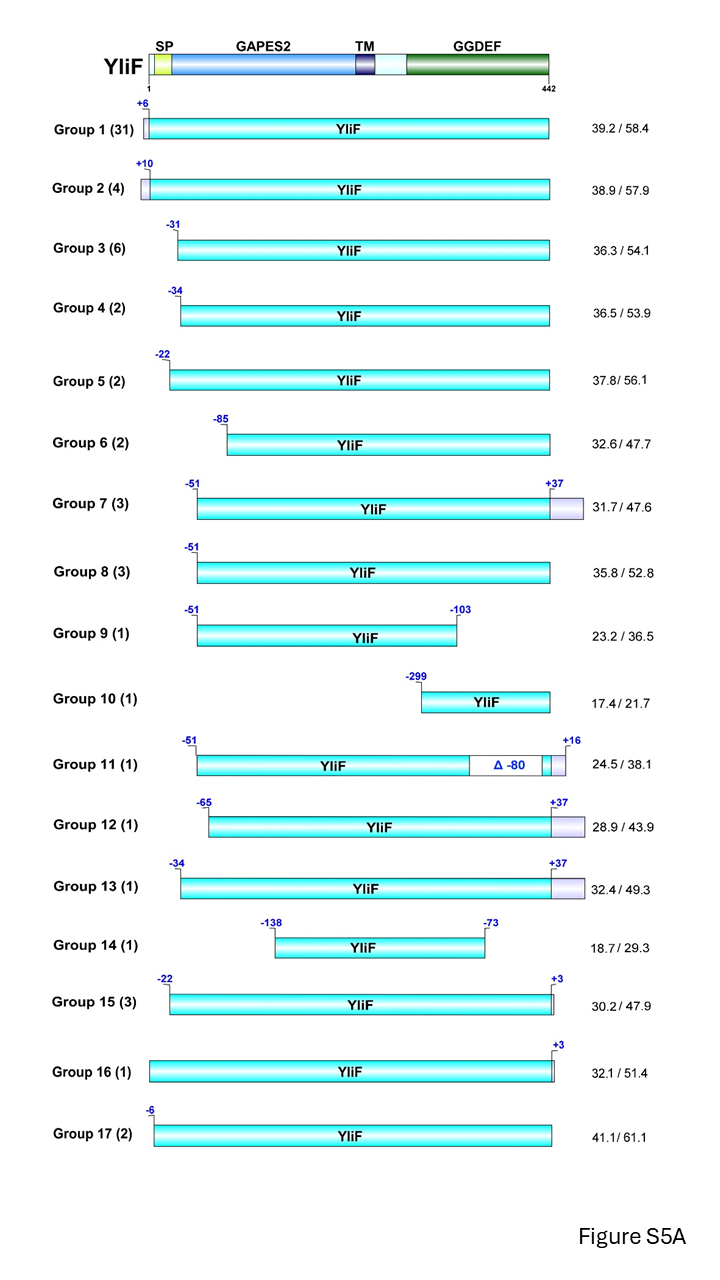

**Figure S5**

Domain structure and length of GGDEF and EAL domain proteins encoded by BcsA encoding *E. coli* plasmids.

**S5A.** YliF GAPES2-GGDEF domain protein structure compared to homologous proteins encoded on BcsA *E. coli* plasmids. On the left, the number of YliF homologous found on plasmids are indicated On the right, the degree of identity and similarity to chromosomally encoded YliF of the reference strain *E. coli* Fec10/MG1655 is indicated.

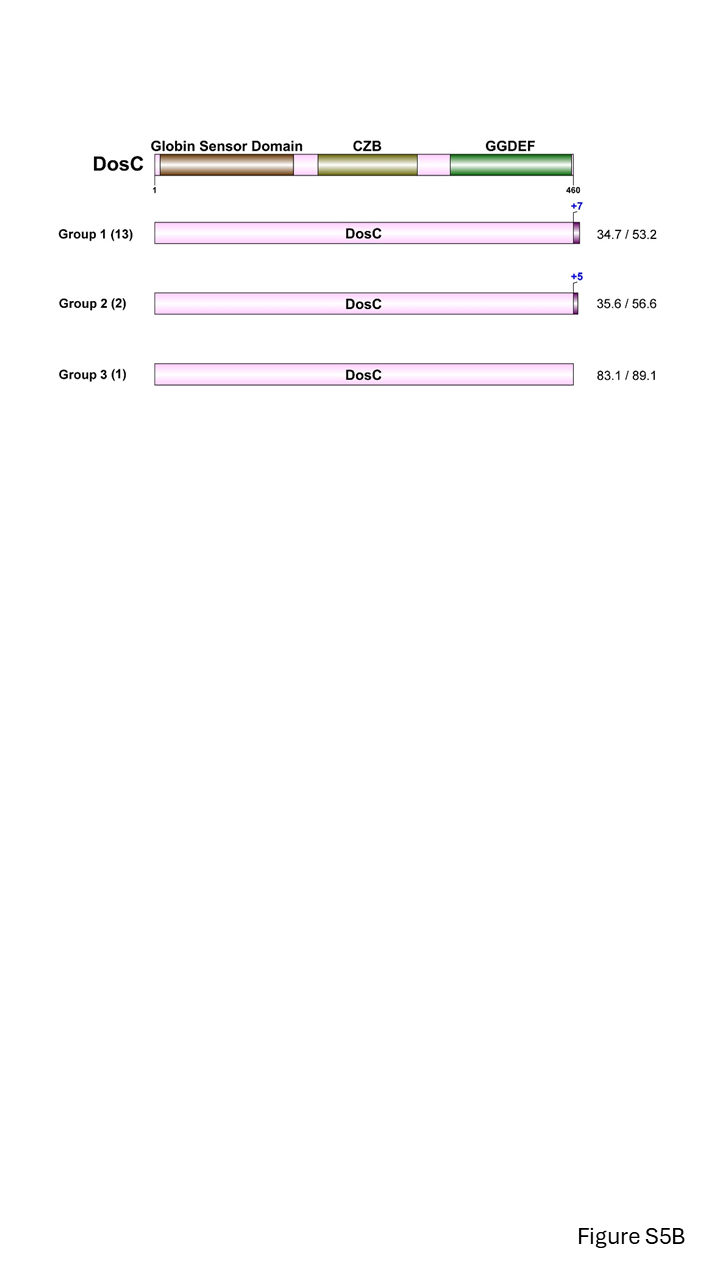

**S5B.** DosC Globin-Sensor-Domain-CZB-GGDEF domain protein structure compared to homologous proteins encoded on BcsA *E. coli* plasmids. On the left, the number of DosC homologous found on plasmids are indicated. On the right, the degree of identity and similarity to chromosomally encoded DosC of the reference strain *E. coli* Fec10/MG1655 is indicated.

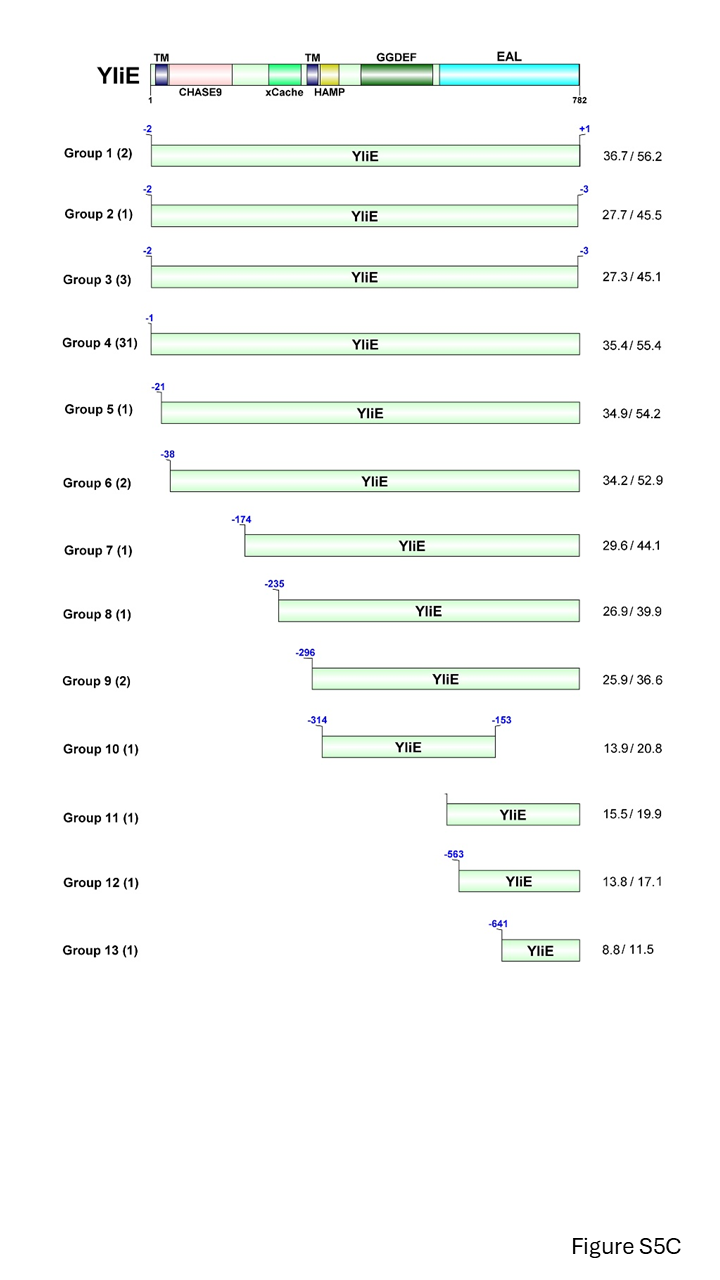

**S5C.** YliE -TM-CHASE9-xCache-TM-HAMP-GGDEF-EAL domain protein structure compared to homologous proteins encoded on BcsA *E. coli* plasmids. On the left, the number of YliE homologous found on plasmids are indicated. On the right, the degree of identity and similarity to chromosomally encoded YliE of the reference strain *E. coli* Fec10/MG1655 is indicated.

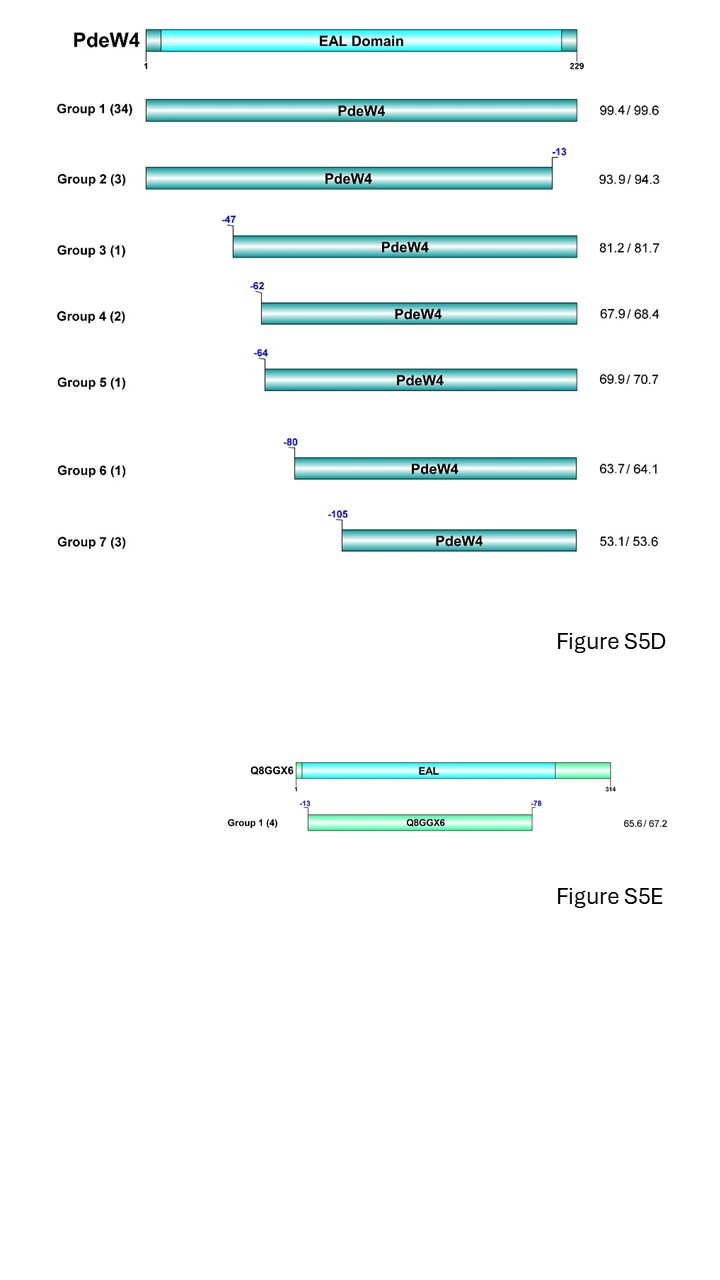

**S5D.** PdeW4 EAL domain protein structure compared to homologous proteins encoded on BcsA *E. coli* plasmids. On the left, the number of PdeW4 homologous found on plasmids are indicated. On the right, the degree of identity and similarity to chromosomally encoded PdeW4 is indicated.

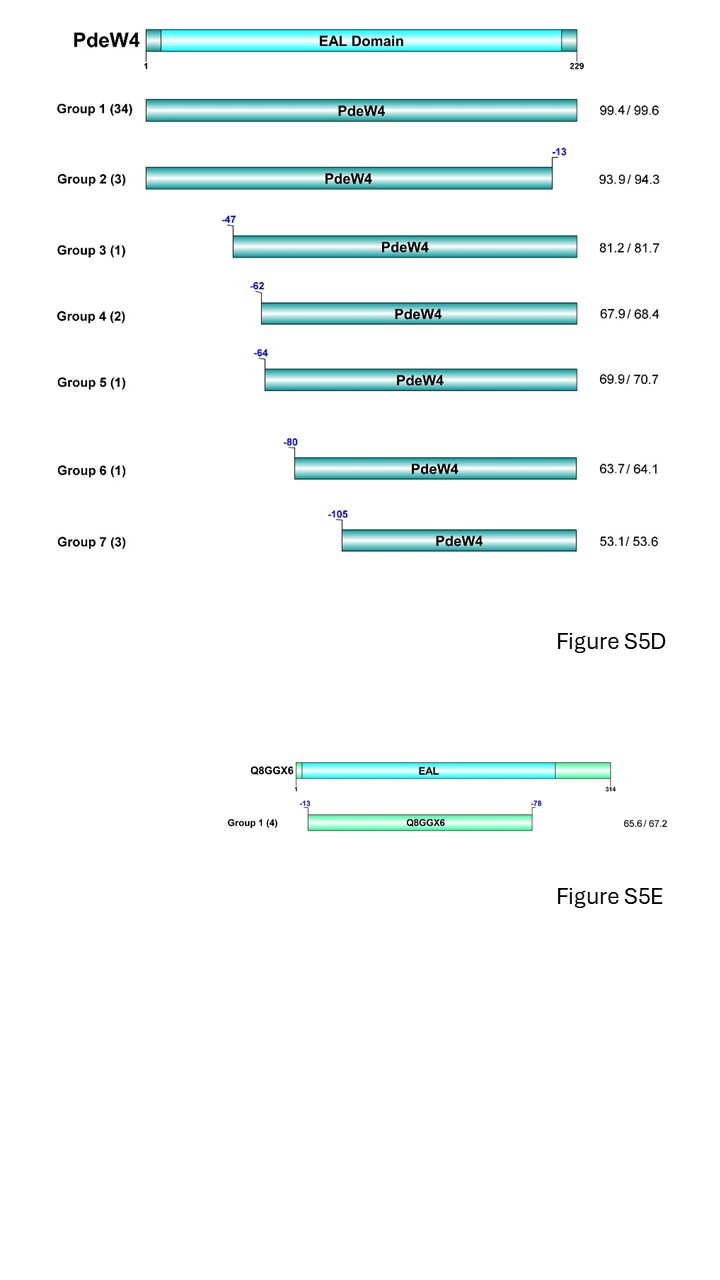

**S5E.** Q8GGX6 EAL domain protein structure compared to homologous proteins encoded on BcsA *E. coli* plasmids. On the left, the number of Q8GGX6 homologs found on plasmids are indicated. On the right, the degree of identity and similarity to chromosomally encoded Q8GGX6 is indicated.

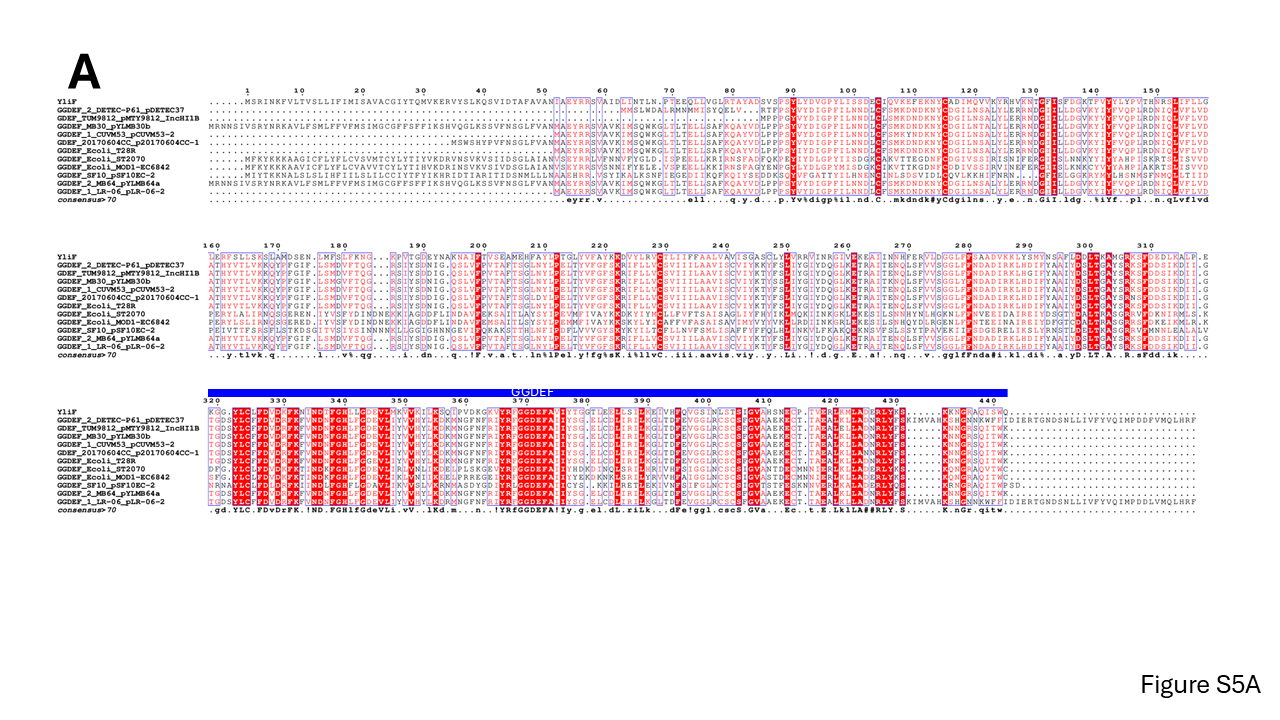

**Figure S6**

**S6A.** Alignment of representative YliF GGDEF domain protein encoded on BcsA bearing plasmids from *E. coli* with the chromosomally encoded YliF reference protein of the reference strain *E. coli* Fec10/MG1655*.* The GGDEF domain with the characteristic GGDEF motif is indicated by a blue bar.

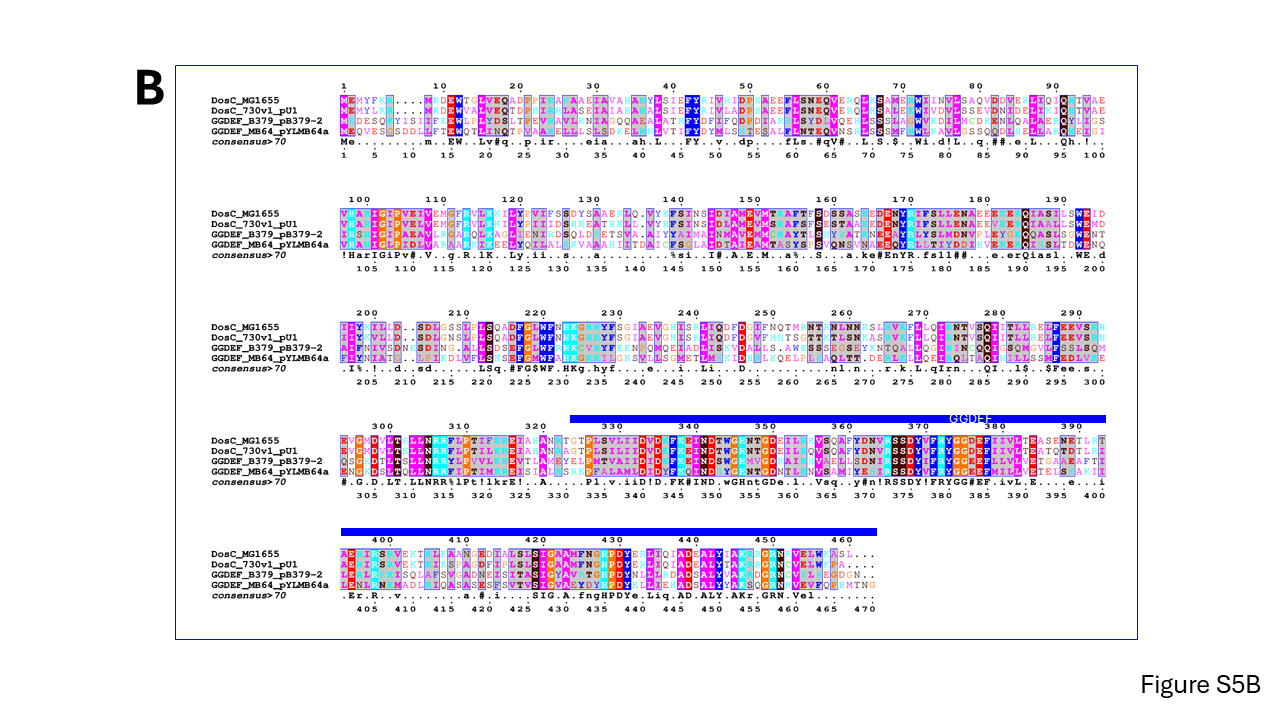

**S6B.** Alignment of representative DosC GGDEF domain protein encoded on BcsA plasmids from *E. coli* with the chromosomally encoded DosC reference protein of the reference strain *E. coli* Fec10/MG1655*.* The GGDEF domain with the characteristic GGDEF motif is indicated by a blue bar.

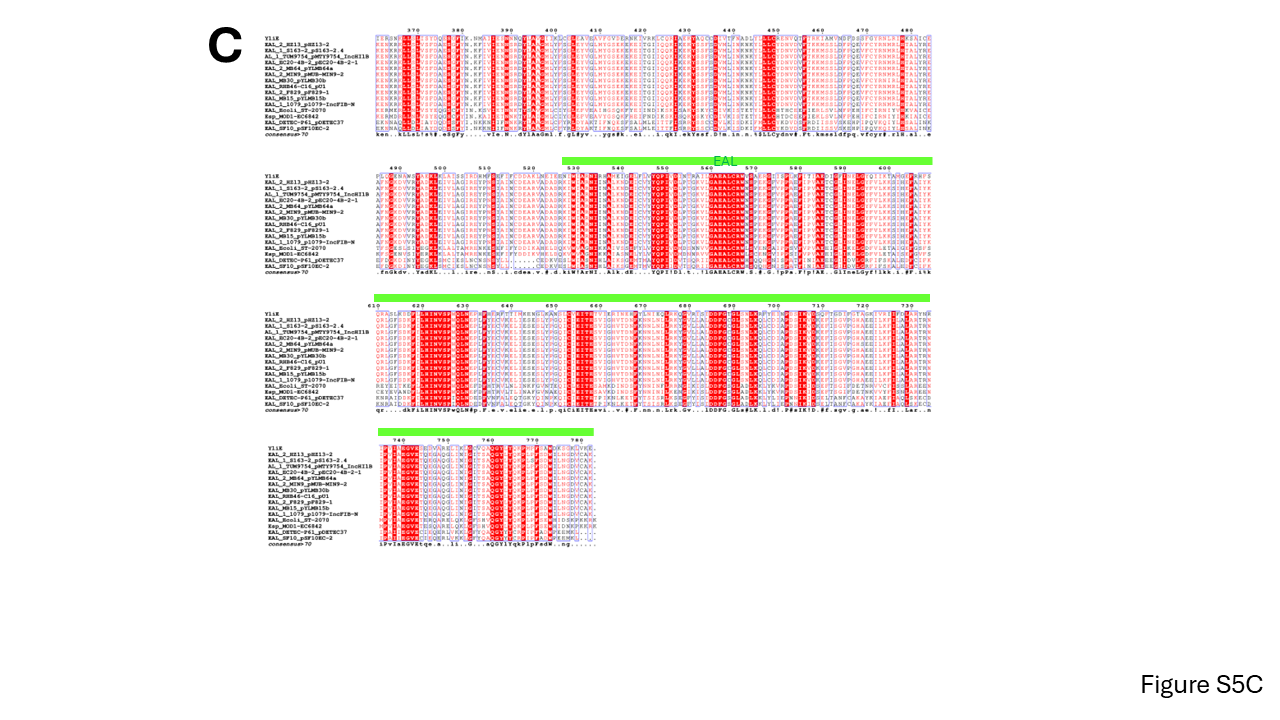

**S6C.** Alignment of representative YliE EAL domain protein encoded on BcsA plasmids from *E. coli* with the chromosomally encoded YliE reference protein of the reference strain *E. coli* Fec10/MG1655*.* The EAL domain with the characteristic EAL motif is indicated by a green bar.

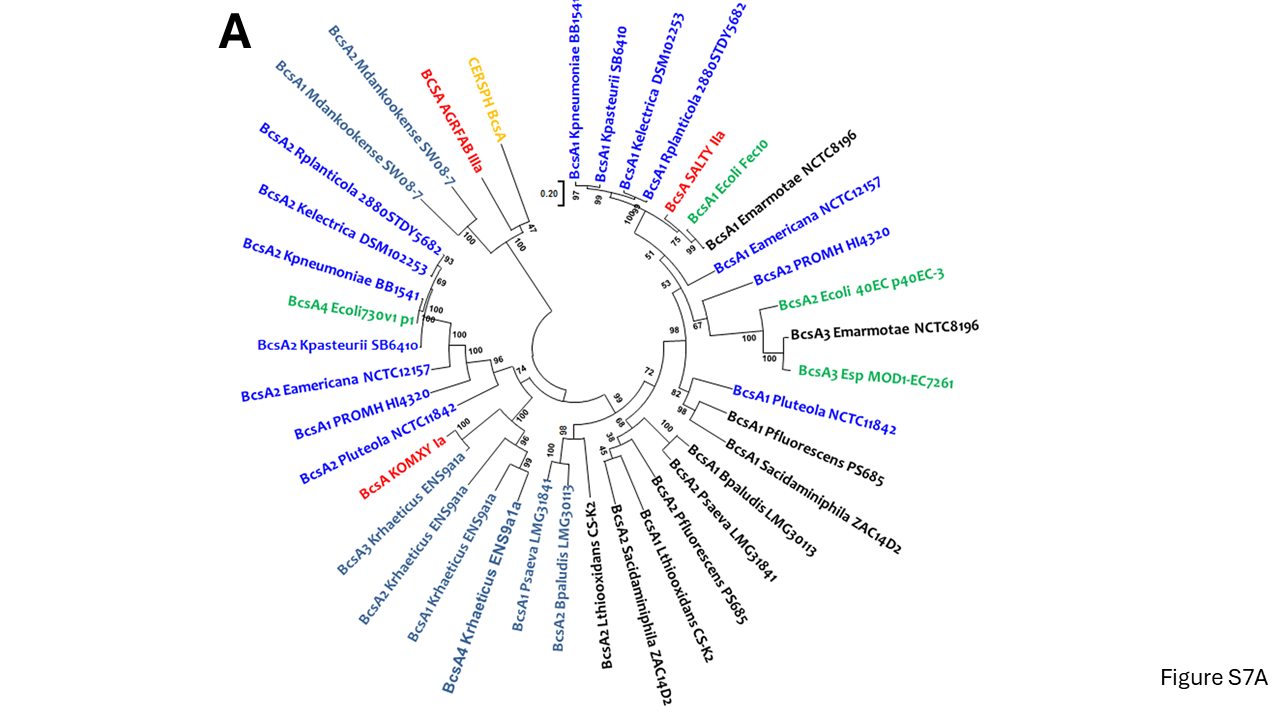

**Figure S7**

**S7A.** Phylogenetic relationship of paralogous and xenologous BcsA cellulose synthases compared to core genome cellulose synthases encoded by individual isolates of a species. BcsA cellulose synthases were retrieved from Uniprot (Apweiler et al. 2004). BcsA cellulose synthases were aligned with ClustalX 2.1 and subsequently manually curated with GeneDoc. The phylogenetic relationship has been assessed using a maximum likelihood (ML) based phylogenetic tree with 1000 bootstraps in MEGA 7.0. Protein identities are; BcsA1_Ecoli_MG1655, P37653; BcsA1_Emarmotae_NCTC8196, WP_000025929.1; BcsA3_Emarmotae_NCTC8196, WP_000576412.1; BcsA1_Kpneumoniae_BB1541, WP_002921508.1; BcsA2_Kpneumoniae_BB1541, WP_004186066.1; BcsA1_Kpasteurii_SB6410, WP_142445849.1; BcsA2_Kpasteurii_SB6410, WP_004126352.1; BcsA1_Kelectrica_DSM102253, WP_141963125.1; BcsA2_Kelectrica_DSM102253, WP_141963111.1; BcsA1_Rplanticola_2880STDY5682; WP_064385327.1; BcsA2_Rplanticola_2880STDY5682, WP_032690497.1; BcsA1_Eamericana_NCTC12157, WP_034790467.1; BcsA2_Eamericana_NCTC12157, WP_034794142.1; BcsA1_Pfluorescens_PS685, WP_150628339.1; BcsA2_Pfluorescens_PS685, WP_150628066.1; BcsA1_Pluteola_NCTC11842, WP_010798019.1; BcsA2_Pluteola_NCTC11842, WP_010795072.1; BcsA1_Sacidaminiphila_ZAC14D2, A0A0S1B309_9GAMM; BcsA2_Sacidaminiphila_ZAC14D2, A0A0S1B2Y2_9GAMM; BcsA1_PROMH_HI4320, B4F0R1_PROMH; BcsA2_PROMH_HI4320, B4F1A9_PROMH; BcsA1_Mdankookense_SW08-7, WP_373321913.1; BcsA2_Mdankookense_SW08-7, VUF12718.1; BcsA1_Bpaludis_LMG30113, WP_031403114.1; BcsA2_Bpaludis_LMG30113, WP_034198152.1; BcsA1_Psaeva_LMG31841, CAG4896730.1; BcsA2_Psaeva_LMG31841, CAG4888940.1; BcsA1_Lthiooxidans_CS-K2, WP_338284421.1; BcsA2_Lthiooxidans_CS-K2, WP_130557523.1; BcsA1_Krhaeticus_ENS9a1a, QIP35774.1; BcsA2_Krhaeticus_ENS9a1a, QIP35696.1; BcsA3_Krhaeticus_ENS9a1a, QIP34621.1; BcsA4_Krhaeticus_ENS9a1a, QIP34889.1; BcsA3_E_coli_MOD1-EC7261, WP_309491259.1; BcsA2_Ecoli_40EC p40EC-3, WP_052248696.1; BcsA4_Ecoli 730V1 p1, WP_004186066.1; CERSPH_BcsA,Q3J125_RHOS4; BcsA_KOMXY_Ia, AHI24410.1; BcsA_SALTY_IIa, A0A0F6B898_SALT1; BCSA_AGRFAB_IIIa, A9CEZ7_AGRFC. Scale bar indicates substitutions per site.

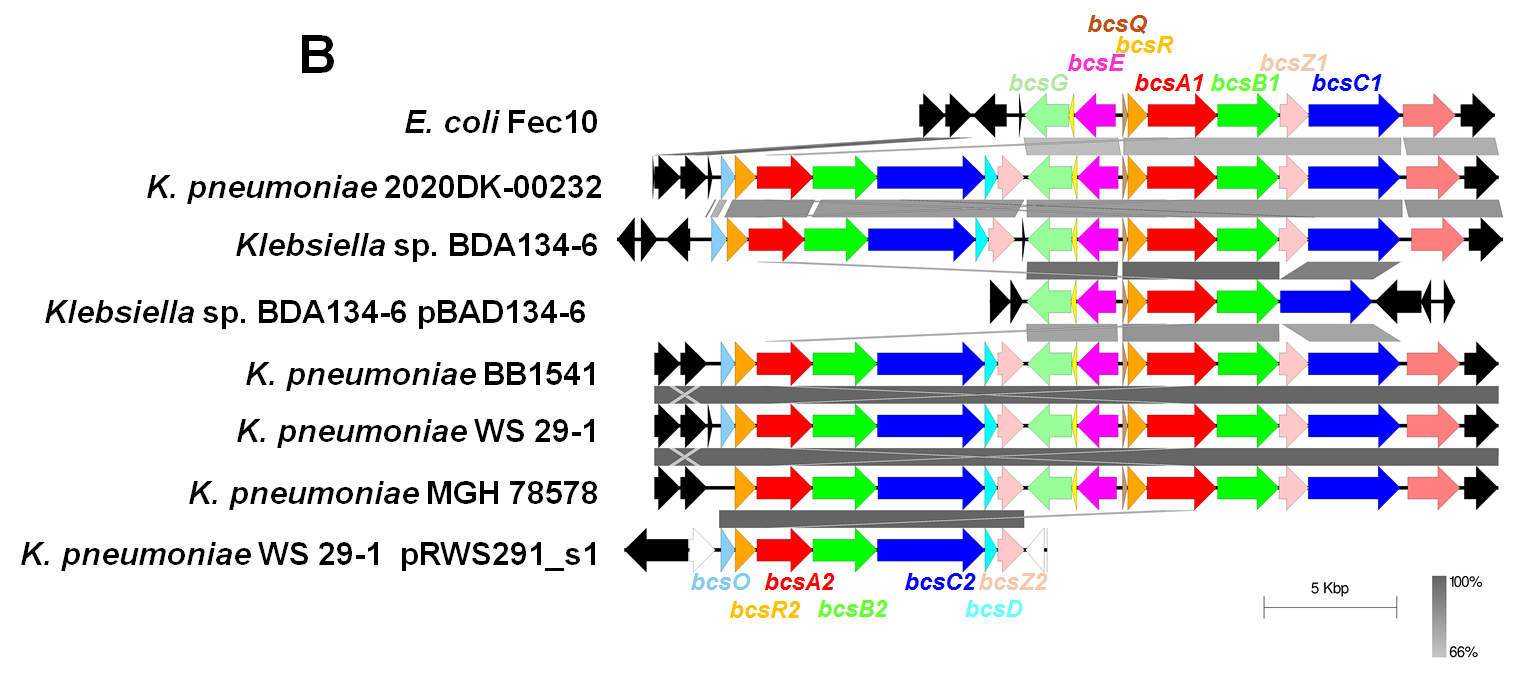

**S7B**. Arrangement of the two cellulose biosynthesis gene clusters in selected *K. pneumoniae* strains. Blast analysis and visualization has been performed with Easyfig after retrieval of the genome annotations of the respective whole genome sequenced strains from NCBI. The *bcs* gene cluster from *E. coli* Fec10 is used as a reference

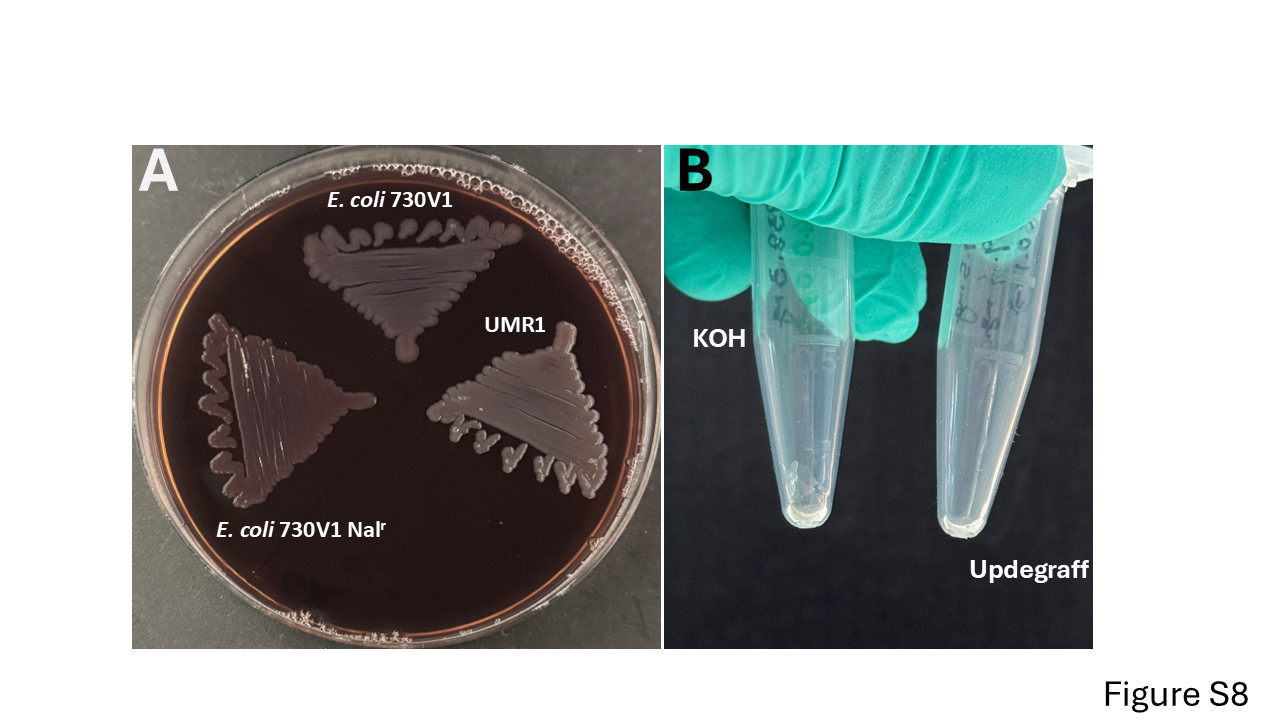

**Figure S8**

**S8A**. Rdar morphotype of *E. coli* 730V1 Nal^r^ compared to the parent strain *E. coli* 730V1 and *S. typhimurium* UMR1. Bacterial cells were grown on a Congo red agar plate incubated for 24 h at 37 ℃ after which the plate was photographed.

**S8B.** Pellet of exopolysaccharide (cellulose) isolated by 1% KOH and the Updegraff approach from *E. coli* 730V1.

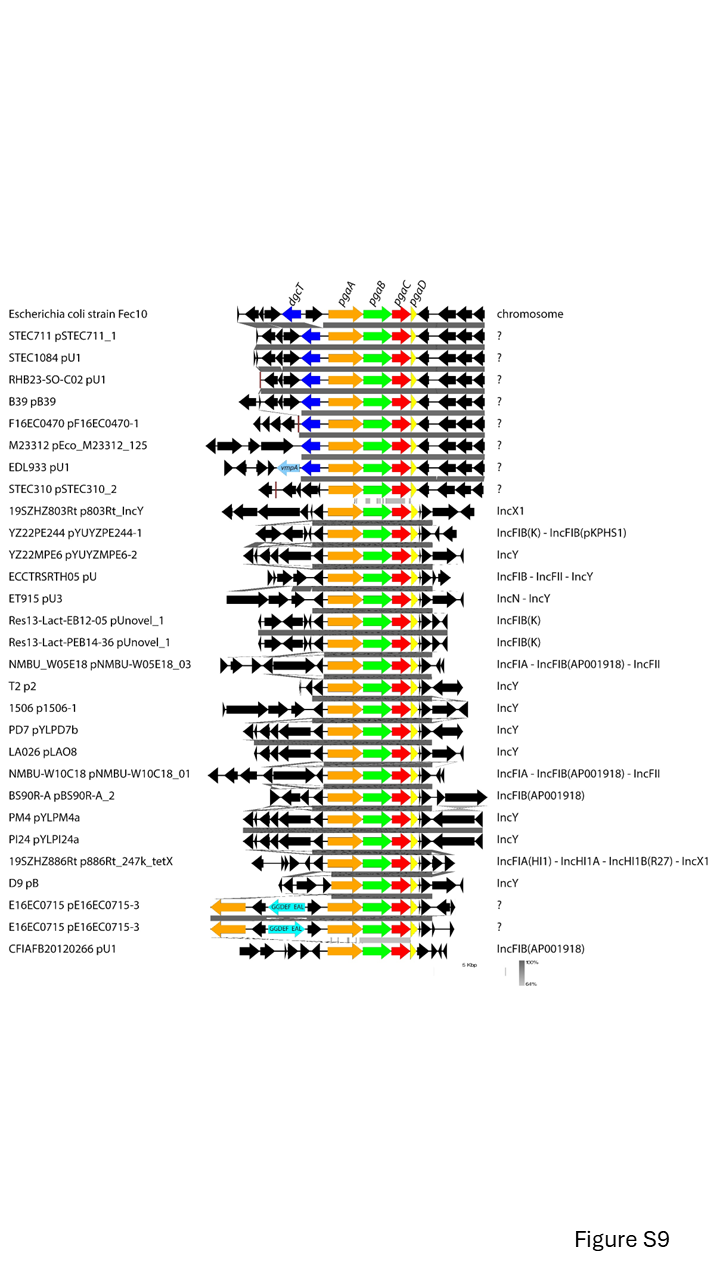

**Figure S9**

Genomic context of mobile element encoded poly-N-acetyl-glucosamine *PgaABCD* biosynthesis gene clusters in comparison with the chromosomally encoded reference *PagABCD* gene cluster. Blue arrows, cyclic di-GMP turnover genes.
